# Outward-oriented sites within clustered CTCF boundaries are key for intraTAD chromatin interactions and gene regulation

**DOI:** 10.1101/2023.05.06.539675

**Authors:** Xiao Ge, Haiyan Huang, Keqi Han, Wangjie Xu, Zhaoxia Wang, Qiang Wu

**Author notes:** These authors contributed equally to this work (Xiao Ge, Haiyan Huang).

## Abstract

CTCF plays an important role in adjusting insulation at TAD boundaries, where clustered CBS (CTCF-binding site) elements are often arranged in a somewhat tandem array with a complex divergent or convergent orientation. Here using *cPcdh* and *HOXD* loci as a paradigm, we look into the clustered CTCF TAD boundaries and find that, counterintuitively, outward-oriented CBS elements are crucial for inward enhancer-promoter interactions as well as for gene regulation. Specifically, by combinatorial deletions of a series of putative enhancer elements *in vivo* or CBS elements *in vitro*, in conjunction with chromosome conformation capture and RNA-seq analyses, we show that deletions of outward-oriented CBS elements weaken the strength of intraTAD promoter-enhancer interactions and enhancer activation of target genes. Our data highlight the crucial role of outward-oriented CBS elements within the clustered CTCF TAD boundaries and have interesting implications on the organization principles of clustered CTCF sites within TAD boundaries.

## Introduction

The interphase genome is organized into highly dynamic structures including chromosome territory, compartment, topologically associated domain (TAD), and chromatin loop^1^. CTCF (CCCTC-binding factor) is a key architecture protein for interphase 3D genome organization and CBS (CTCF binding site) elements throughout the mammalian genome function as insulators to block aberrant enhancer activation and ensure proper gene expression or to mark the boundaries between euchromatin and heterochromatin^2-4^. A prevailing model for interphase chromosome folding, known as loop extrusion, posits that cohesin complex extrudes chromatin fibers bidirectionally into expanding loops until blocked asymmetrically by orientated CBS elements^5-7^. This model explains TAD formation and is consistent with the observations that the two flanking boundaries of each TAD are associated with mostly convergent forward-reverse CBS elements^8,9^. However, it is puzzling that most TAD boundaries contain clustered CBS elements with complex orientations of both convergence and divergence.

TAD boundaries emerged as insulator elements that restrain interTAD chromatin interactions. Their dysfunction leads to abnormal development and progressive oncogenesis^10-12^. Mammalian clustered CTCF TAD boundaries are more evolutionarily conserved than other genomic regions^13-15^. Higher insulation scores of TAD boundaries are associated with rich CTCF enrichments and clustered enhancers known as super-enhancers tend to be insulated by strong boundaries^16,17^. Divergent CBS elements are enriched at TAD boundaries; however, a divergent orientation signature is not strictly required for effective insulation. For example, the characteristics of specific CBS elements can outweigh CBS number and orientation^18^. Finally, insulation potency depends on the local context of CBS elements and robustness of enhancer-promoter communications can bypass CTCF insulation regardless of the intervening CBS strength^19-22^.

As a paradigm for investigating mechanisms of 3D genome folding, the mouse clustered protocadherins (*cPcdh*) comprise three sequential gene clusters, *Pcdh α, β*, and *γ*, encoding 58 isoforms, spanning ∼1 million bps (Fig. 1a)^23^. The *Pcdh α* and *γ* clusters consist of 14 and 22 variable first exons, respectively, each of which is *cis*-spliced to single sets of three downstream constant exons; whereas the *Pcdhβ* cluster consists of 22 single exons (*β1*-*β22*) with no constant exon. The *cPcdh* promoter choice is determined by CTCF/cohesin-mediated long-range enhancer-promoter looping interactions anchored by convergent CBS elements^9,24,25^. Their combinatorial expression patterns in single neurons generate enormous cell-surface molecular diversity for assembling enormous networks of neuronal connectivity^26,27^. In addition, precise expression patterns of members of the *HoxD* gene cluster are central for limb development^28,29^. Here, using *cPcdh* and *HOXD* as model genes, we find that outward-oriented CBS elements within clustered CTCF boundaries are crucial for intraTAD promoter-enhancer interactions and gene regulation. Our data shed interesting lights on the organization principles of the clustered CTCF TAD boundaries.

**Fig. 1.**
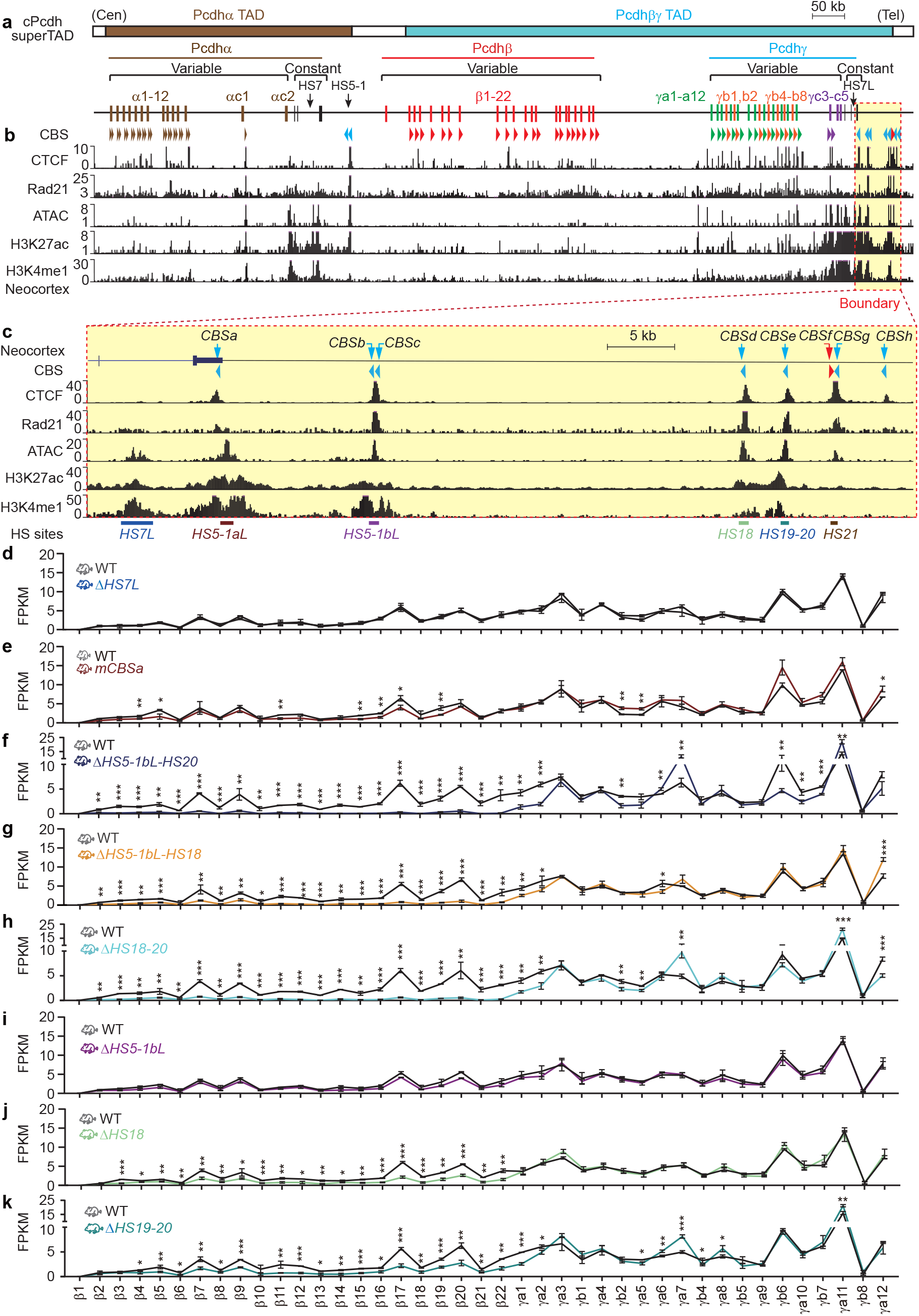
The downstream *cPcdh* superTAD boundary comprises clustered cis-regulatory elements for *Pcdhβγ* gene regulation. a, Schematics of the mouse *cPcdh* locus. The mouse *Pcdh α, β*, and *γ* gene clusters are closely-linked within the *cPcdh* superTAD, which is divided into two TADs of *Pcdhα* and *Pcdhβγ. Pcdh α* and *γ* gene clusters contain 14 and 22 variable first exons, respectively, each of which is alternatively spliced to a single set of three downstream constant exons. The *Pcdhβ* gene cluster contains 22 variable exons but with no constant exon. Two *Pcdhα* enhancers, *HS5-1* and *HS7*, are indicated by black arrows. **b**, ChIP-seq of CTCF, Rad21, H3K27ac, and H3K4me1 as well as ATAC-seq profiles of the *cPcdh* locus in the mouse neocortex. Arrowheads indicate CBS elements with orientations. Within the *Pcdhα* TAD, each *Pcdhα* alternate promoter is flanked by two forward CBS elements, the downstream enhancer of *HS5-1* is flanked by two reverse CBS elements. Within the *Pcdhβγ* TAD, each *Pcdh β* or *γ* promoter is associated with one forward CBS, except for *β1, γc4*, and *γc5* which have no CBS, as well as *γc3* which has two forward CBS elements. **c**, Close-up of the *cPcdh* superTAD downstream boundary marked by a red dotted rectangle in **a** and **b**. The boundary comprises eight CBS elements, of which only *CBSf* is in the reverse orientation, and six ATAC-seq peaks, of which *HS7L, HS5-1bL, HS18*, and *HS19-20* are knocked out individually or in combinations in mice. *HS5-1aL* is mutated at the *CBSa* motif (*mCBSa*) due to its location in the coding region. **d-k**, Expression levels of the *Pcdh β* and *γ* genes in the mouse neocortex from Δ*HS7L, mCBSa*, Δ*HS5-1bL-HS20*, Δ*HS5-1bL-HS18*, Δ*HS18-20*, Δ*HS5-1bL*, Δ*HS18*, or Δ*HS19-20* homozygous mice compared to their wild-type (WT) littermates. FPKM, fragments per kilobase of exon per million fragments mapped. Data as mean ± SD. *P < 0.05, **P < 0.01, ***P < 0.001.

## Results

### The downstream boundary of *cPcdh* superTAD comprises a cluster of CTCF sites with mixed orientations

During mouse neocortical development, the 58 *cPcdh* genes are organized into a large megabase-sized superTAD comprised of the *Pcdhα* and *Pcdhβγ* TADs (Fig. 1a; Supplementary Fig. 1a-c)^9,30^. At the telomeric boundary of the *cPcdh* superTAD, there is a complex array of CTCF sites or CBS elements (*CBSa-h*, Fig. 1a-c). Among these eight CBS elements, *CBSa-e, CBSg*, and *CBSh* are in the reverse orientation (Supplementary Fig. 1d). By contrast, a single *CBSf* is in the forward orientation (Supplementary Fig. 1e). We performed chromatin immunoprecipitation followed by high-throughput sequencing (ChIP-seq) experiments with a specific antibody against CTCF using microdissected mouse neocortical tissues and found that CTCF is enriched at these elements (Fig. 1c). Interestingly, the *CBS b* and *c* elements form a single peak and *CBS f* and *g* elements form a single peak (Fig. 1c). To see whether these elements are anchors of cohesin loop extrusion, we performed ChIP-seq experiments with a specific antibody against Rad21, a subunit of cohesin, and found that cohesin is co-localized with CTCF at all of these CBS elements, suggesting that cohesin loop extrusion from this boundary may regulate *cPcdh* gene expression (Fig. 1b,c).

We first comprehensively mapped expression patterns of each member of the three *Pcdh* gene clusters at a series of 12 developmental stages by collecting embryos every other day and new pubs every other two days. RNA-seq of microdissected neocortical tissues showed that each member of the *cPcdh* genes is expressed dynamically and that the *cPcdh* genes are most abundantly expressed at the newborn stage (Extended Data Fig. 1). Thus, we used the newborn mouse neocortical tissues in the following gene regulation studies.

Within the *Pcdhα* TAD, members of the *Pcdhα* cluster are activated by *HS5-1* and *HS7* (Fig. 1b and Extended Data Fig. 2a,b)^24,31,32^. However, how members of the *Pcdhβγ* clusters are regulated within the *Pcdhβγ* TAD is not clear. To this end, we profiled the chromatin regulatory landscape by assaying for transposase-accessible chromatin with high-throughput sequencing (ATAC-seq) and identified six highly accessible regions at the telomeric boundary of the *Pcdhβγ* TAD, corresponding to the previously identified DNaseI hypersensitive sites of *HS7L, HS5-1aL, HS5-1bL, HS18, HS19-20*, and *HS21* (Fig. 1c and Extended Data Fig. 2c,d)^24,33^. Finally, we performed H3K27ac and H3K4me1 ChIP-seq and found, except the last one (*HS21*), all of the other five clustered *HS* elements are marked by H3K27ac and H3K4me1, suggesting that they form a super-enhancer for members of the *Pcdhβγ* gene clusters (Fig. 1c).

### The *Pcdhβγ* TAD boundary comprises a super-enhancer

We dissected the putative enhancers in mice *in vivo* by CRISPR/Cas9-based DNA-fragment editing system (Extended Data Fig. 3a-c)^9^ to test their functionality. We first deleted the *HS7L* element (Extended Data Fig. 3b,c) and found, surprisingly, it does not lead to decreased levels of *cPcdh* gene expression in neocortical and olfactory tissues (Fig. 1d, Extended Data Fig. 3e-g, and Supplementary Fig. 2a). Because *HS5-1aL* is in the *Pcdhγ* constant coding region and its deletion will destroy the transcript integration, we mutated the *CBSa* element within *HS5-1aL* by homologous recombination using CRISPR with an ssODN donor. We found that *CBSa* mutation abolished CTCF enrichments (Extended Data Fig. 3d). Interestingly, it only results in a moderate decrease of expression levels of members of the *Pcdhβγ*, but not *α*, clusters (Fig. 1e, Extended Data Fig. 3e, and Supplementary Fig. 2b), suggesting a limited role of *CBSa* in tethering *HS5-1aL* to *Pcdhβγ*.

We next deleted the entire region covering all of the other three putative enhancers (from *HS5-1bL* to *HS19-20*) and found that, in contrast to *HS7* and *HS5-1aL*, this large deletion leads to a significant decrease of levels of *Pcdhβγ* gene expression, suggesting that enhancers for members of *Pcdhβγ* clusters reside in this region (Fig. 1f and Supplementary Fig. 2c). We then dissected this region in great details and found that deletion of the large DNA fragment covering both *HS5-1bL* and *HS18* results in a significant decrease of levels of *Pcdhβγ* gene expression (Fig. 1g and Supplementary Fig. 2d). In addition, deletion of the DNA fragment covering both *HS18* and *HS19-20* also results in a significant decrease of levels of *Pcdhβγ* gene expression (Fig. 1h and Supplementary Fig. 2e).

Finally, we deleted each of these three elements individually and found that, surprisingly, deletion of *HS5-1bL* has no effect on *Pcdhβγ* gene expression (Fig. 1i and Supplementary Fig. 2f). By contrast, deletion of either *HS18* or *HS19-20* result in a significant decrease of levels of *Pcdhβγ* gene expression (Fig. 1j,k and Supplementary Fig. 2g,h). These data suggest that these two H3K27ac-enriched elements play a leading role in activating *Pcdhβγ* genes *in vivo* in the mouse brain. Consistently, 4C experiments revealed close contacts between *Pcdhβγ* and *HS18-20* (Extended Data Fig. 4a,b). Unlike *HS5-1*, which is brain-specific, *HS18-20* are enriched with H3K27ac and H3K4me3 marks in multiple tissues (Extended Data Fig. 4c,d), explaining why members of the *Pcdhβγ* gene clusters are expressed in both neural and non-neural tissues. Taken together, these data suggest that the *Pcdhβγ* downstream TAD boundary is a superenhancer comprising a cluster of enhancers among which *HS18* and *HS19-20* are the most important.

### *HS5-1bL* is a *Pcdhγc3*-specific insulator

*Pcdhγc3* is the most abundantly expressed isoform in the mouse neocortex and has a unique role in dendrite arborization (Extended Data Fig. 1e)^34^. However, the underlying mechanism remains unclear. To this end, we performed 4C experiments and found that *Pcdhγc3* is in close contacts with *HS5-1bL, HS18*, and *HS19-20* (Fig. 2a). Consistently, deletion of *HS18* and/or *HS19-20* results in a significant decrease of levels of *Pcdhγc3* gene expression, suggesting that *HS18* and *HS19-20* are enhancers for *Pcdhγc3* (Fig. 2b-d). Surprisingly, deletion of *HS5-1bL* leads to a significant increase of levels of *Pcdhγc3*, but not *γc4* or *γc5*, gene expression (Fig. 2e), suggesting that *HS5-1bL* is a putative insulator to specifically block the activation of *Pcdhγc3* by the *HS18* and *HS19-20* enhancers. To this end, we performed 4C experiments with *Pcdhγc3* as an anchor using neocortical tissues of the *HS5-1bL* deleted mice and found a significant increase of long-distance chromatin interactions between *Pcdhγc3* and *HS18-20* (Fig. 2f). To confirm this aberrantly increased chromatin contacts upon deletion of *HS5-1bL*, we performed 4C with *HS19-20* as an anchor and indeed observed increased chromatin contacts with *Pcdhγc3* (Fig. 2g). In conjunction with no expression increase of all of the other members of the *Pcdhβγ* gene clusters upon deletion of *HS5-1bL* (Fig. 1i), we conclude that *HS5-1bL* is a *γc3*-specific insulator.

**Fig. 2.**
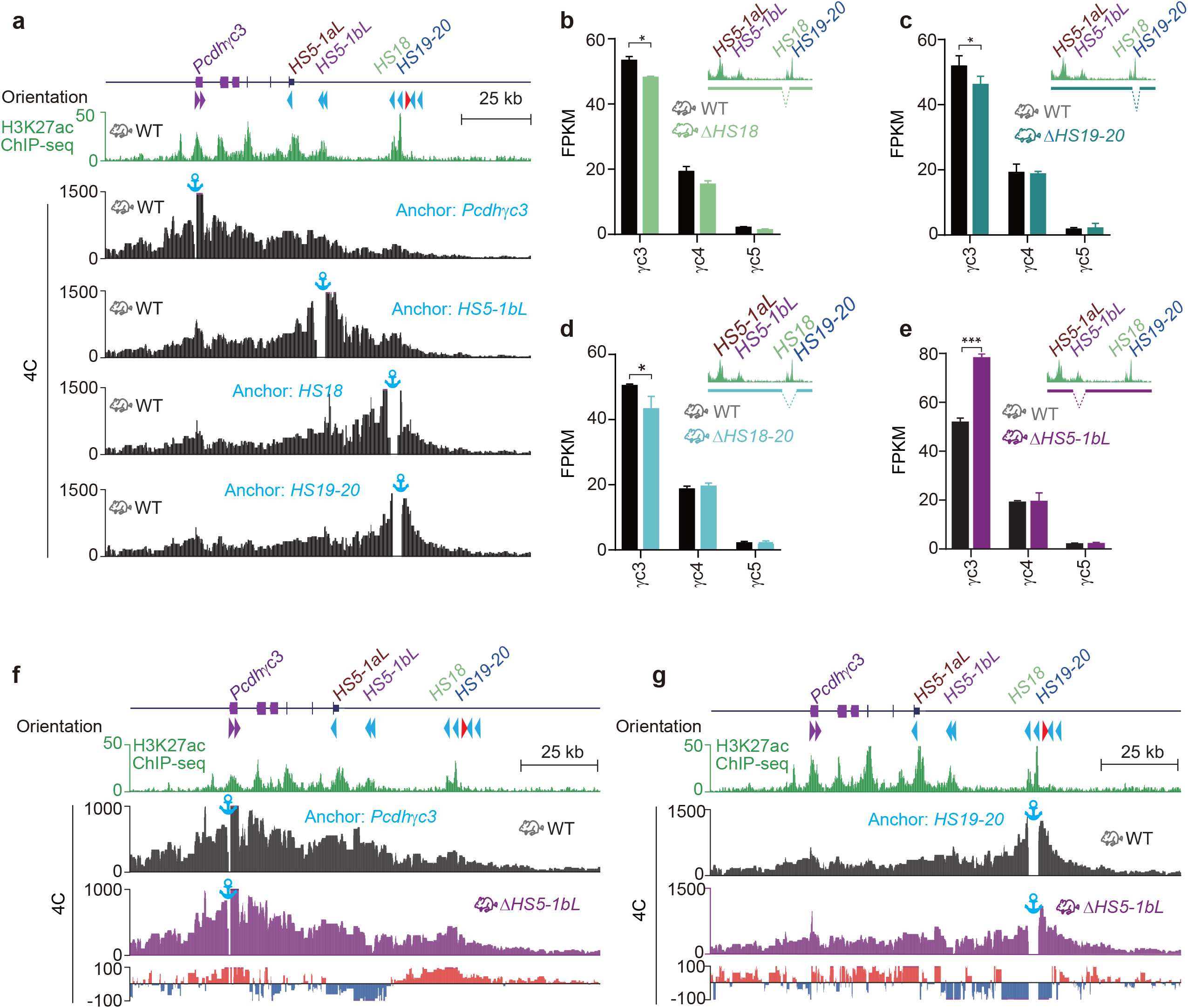
*HS5-1bL* is an insulator specific for *Pcdhγc3*. a, 4C profiles using a repertoire of elements as anchors showing close contacts between *Pcdh*γ*c3* and *HS5-1bL, HS18*, or *HS19-20* in wild-type (WT) mice. **b-e**, Expression levels of the *Pcdhγ* c-type genes in the mouse neocortex from Δ*HS18* (**b**), Δ*HS19-20* (**c**), Δ*HS18-20* (**d**), *or* Δ*HS5-1bL* (**e**) homozygous mice compared to their wild-type littermates. Note the significant increase of expression levels of *Pcdhγc3* upon deletion of *HS5-1bL*. Data as mean ± SD. *P < 0.05, **P < 0.01, ***P < 0.001. **f**, 4C profiles using the *Pcdhγc3* promoter as an anchor showing increased chromatin interactions between *Pcdhγc3* and its *HS18-20* enhancers in Δ*HS5-1bL* homozygous mice compared to their WT littermates. Differences (Δ*HS5-1bL* versus WT) are shown under the 4C profiles. **g**, 4C profiles using *HS19-20* as an anchor showing increased chromatin interactions between *Pcdhγc3* and *HS19-20* in Δ*HS5-1bL* mice compared to their WT littermates. Differences (Δ*HS5-1bL* versus WT) are shown under the 4C profiles.

### *cPcdhs* are expressed from H3K9me3-enriched facultative heterochromatins of superTAD

We next mapped the chromatin landscape of a wide variety of histone modifications in the *cPcdh* superTAD and found that this region is enriched for both repressive marks of H3K9me3 and active marks of H3K36me3 (Fig. 3a), suggesting that *cPcdhs* are expressed from facultative heterochromatin region of the superTAD. In addition, expression of each member of the three *Pcdh* gene clusters is strongly correlated with active marks of H3K4me3 and H3K9ac (Fig. 3b and Supplementary Figs. 3,4). Moreover, this region is also marked by the repressive histone marks of H3K9me2 in the mouse neocortex (Fig. 3c and Extended Data Fig. 5). Specifically, the coding region of each member of the *Pcdh* clusters is correlated with a peak of H3K9me2 (Extended Data Fig. 5) but not H3K9me3 (Extended Data Fig. 6). To further investigate mechanisms of *cPcdh* gene expression, we performed methylated DNA immunoprecipitation sequencing (MeDIP-seq) experiments and found that each member of the H3K9me3-enriched *cPcdhs* is hypermethylated on the body of each variable exon (Fig. 3c, Extended Data Fig. 7a-d). We noted that *Pcdhβ1* differs from other *cPcdh* members in that its DNA-methylation peak is much closer to the transcription start site (Extended Data Fig. 7e), explaining why it is silenced in the mouse neocortex (Fig. 3b).

**Fig. 3.**
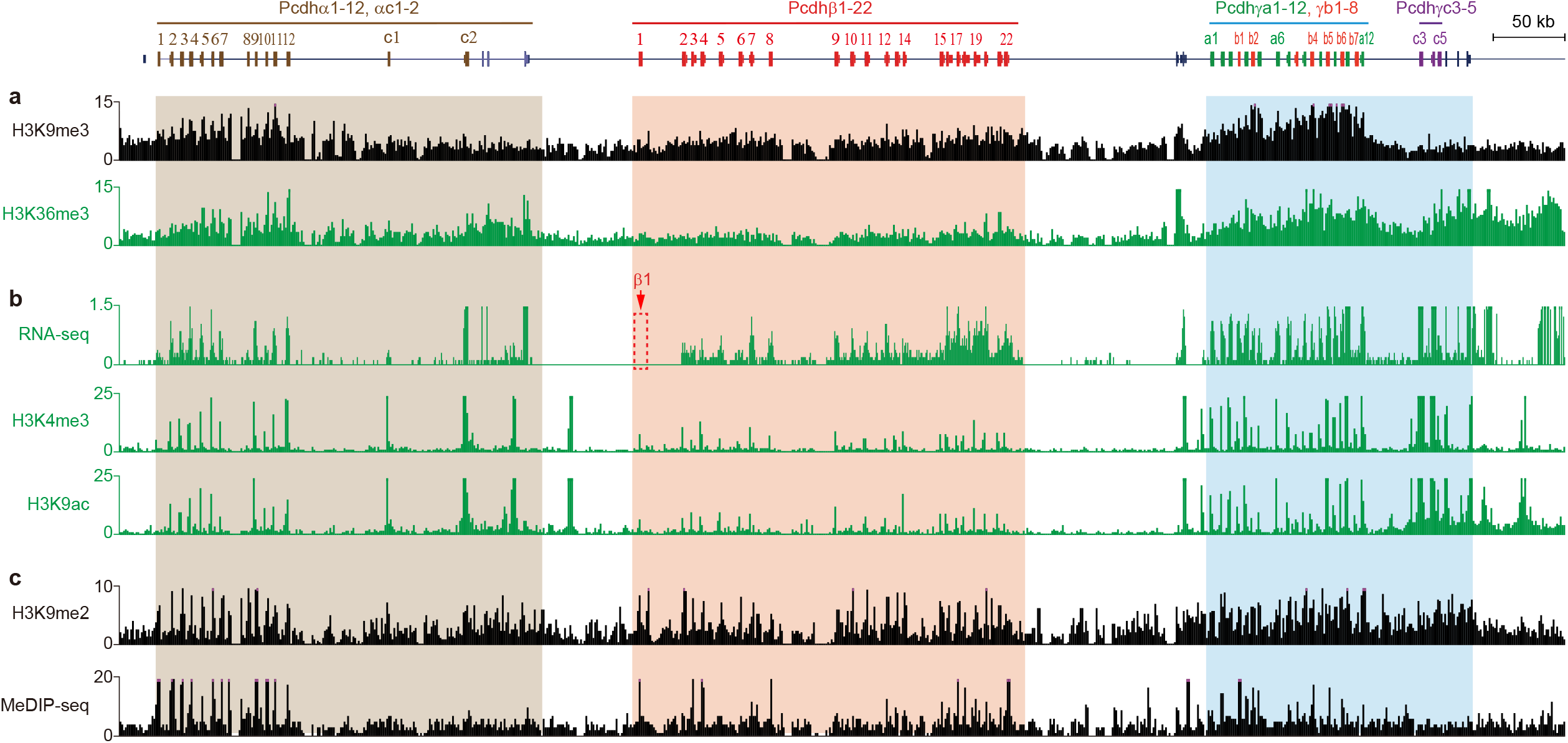
The *cPcdh* genes are expressed from H3K9me3/H3K36me3-enriched facultative heterochromatin domains. **a**, H3K9me3 and H3K36me3 ChIP-seq profiles of the three *Pcdh* clusters in mouse neocortex showing colocalization of both the active mark of H3K36me3 and inactive mark of H3K9me3. **b**, RNA-seq as well as H3K4me3 and H3K9ac ChIP-seq profiles of the *cPcdh* clusters in mouse neocortex showing a strong correlation between each c*Pcdh* gene and the enrichment of active marks. **c**, The H3K9me2 ChIP-seq and MeDIP-seq profiles of the *cPcdh* clusters showing enrichments of the repressive mark of H3K9me2 and DNA methylation at most *cPcdh* genes in the mouse neocortex.

### Outward-oriented *CBSf* is the key boundary CTCF site

All of the *HS5-1aL, HS5-1bL, HS18*, and *HS19-20* enhancers contain reverse-oriented CBS elements (*CBSa-e*, Fig. 1c) that are enriched for cohesin and are in close contacts with the forward-oriented CBS elements of the *Pcdhβγ* promoters via CTCF/cohesin-mediated chromatin loops. In addition, there are two more reverse-oriented CBS elements, *CBSg* and *CBSh*, located immediately downstream of this super-enhancer. However, there is a single forward-oriented *CBSf* element located between the super-enhancer and the two reverse-oriented *CBS g* and *h* elements (Fig. 4a). To this end, we generated a *CBSf*-deletion mouse line (Fig. 4a and Extended Data 8a) and confirmed the abolishment of CTCF and cohesin enrichments at the location of *CBSf* but not *CBSg* (Fig. 4a,b).

**Fig. 4.**
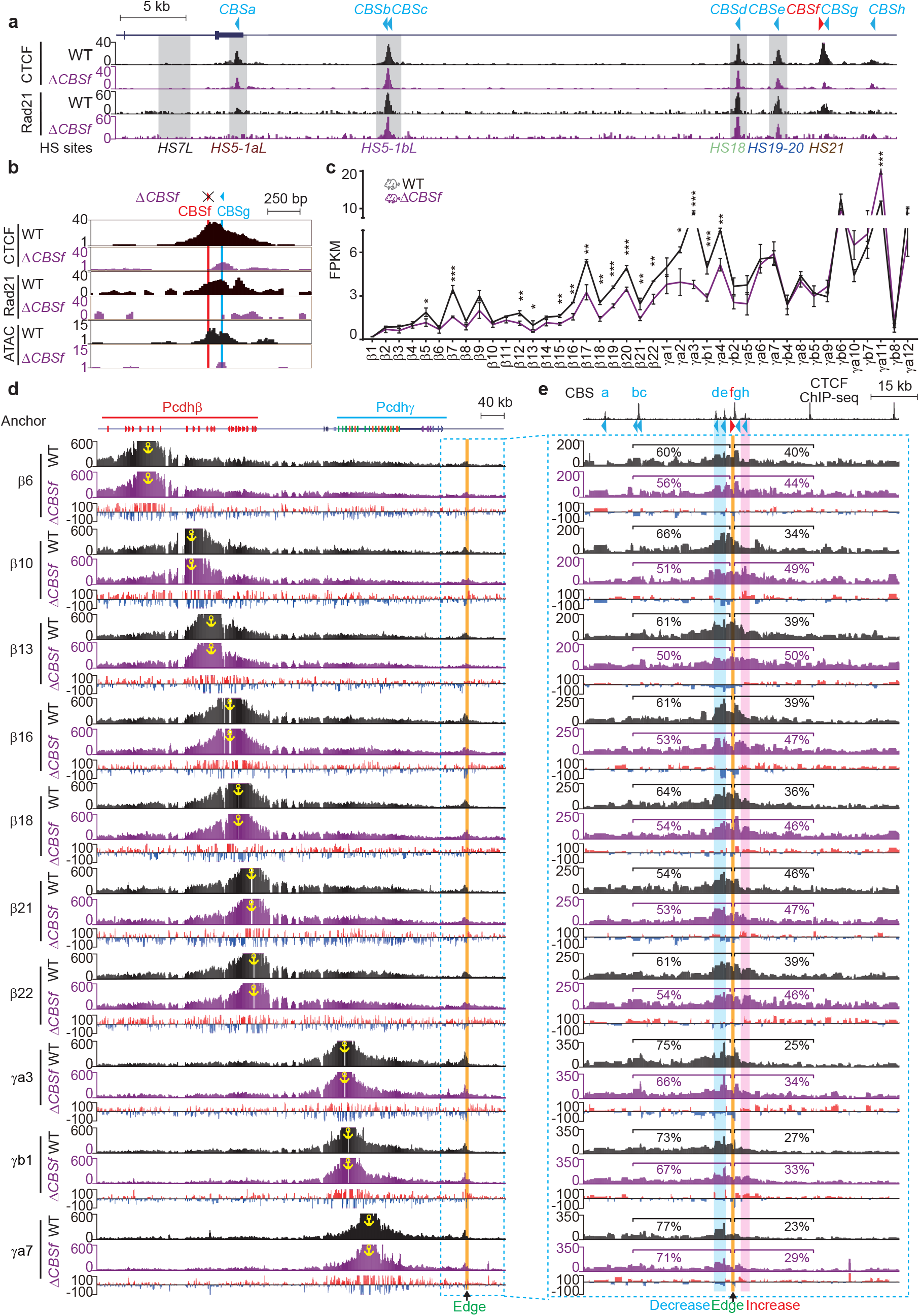
Outward-oriented *CBSf* regulates *Pcdhβγ* by blocking their aberrant looping outside of the TAD boundary. **a**, CTCF and Rad21 ChIP-seq profiles of the *Pcdh****βγ*** downstream TAD boundary in *CBSf-*deleted (Δ*CBSf*) homozygous mice compared to their wild-type (WT) littermates. **b**, Close-up of CTCF and Rad21 ChIP-seq as well as ATAC-seq profiles in Δ*CBSf* mice compared to their WT littermates. **c**, RNA-seq showing decreased expression levels of *Pcdhβγ* upon *CBSf* deletion. Data as mean ± SD, \**p* < 0.05, \*\**p* < 0.01, \*\*\**p* < 0.001; one-tailed Student’s *t* test. **d-e**, 4C profiles using a repertoire of *Pcdhβγ* promoters as anchors showing decreased chromatin interactions with *HS18-20* enhancers (highlighted in blue, **e**) and increased chromatin interactions beyond *CBSf* (highlighted in pink, **e**). Note a sharp edge for the transition from decrease to increase at the location of *CBSf*. Differences (Δ*CBSf* versus WT) are shown under the 4C profiles.

Remarkably, deletion of the forward-oriented *CBSf* element results in a significant decrease of expression levels of members of the *Pcdhβγ* gene clusters (Fig. 4c and Supplementary Fig. 5a), demonstrating that the single forward-oriented *CBSf* plays a crucial role in the regulation of *Pcdhβγ* gene expression. To investigate the underlying mechanism, we performed a series of chromatin conformation capture experiments with members of the *Pcdhβγ* gene clusters as anchors and found a significant decrease of long-distance chromatin contacts between the super-enhancer and members of the *Pcdhβγ* gene clusters (Fig. 4d). Finally, we observed a significant increase of long-distance chromatin contacts with *CBSh* with a sharp edge at the location of *CBSf* (Fig. 4e). To confirm this, we performed reciprocal capture experiments with *CBSh* as an anchor and observed a significant increase of long-distance chromatin contacts with members of the *Pcdhβγ* gene clusters (Extended Data Fig. 8b). Finally, combined deletion of both *CBSf* and *CBSg* elements (Extended Data Fig. 8c) results in a similar phenotype (Extended Data Figs. 8d,9 and Supplementary Fig. 5b). Taken together, we conclude that a single outward-oriented *CBSf* element within the clustered CTCF TAD boundary plays a key role in *Pcdhβγ* gene regulation.

### Outward-oriented CBS elements within a clustered CTCF TAD boundary are crucial for *HOXD13* expression

To see whether it is a general phenomenon for a key role of the outward-oriented boundary CBS element in gene regulation, we investigated the *HOXD* left boundary, which has a complex array of six conserved clustered CBS elements (*CBS1-6*, Supplementary Fig. 6a-d), of the human centromeric TAD (C-DOM) (Fig. 5a-b)^28,29^. The human *HOXD* locus is located at the right boundary of the C-DOM (Extended Data Fig. 10a,b). It contains nine *HOXD* genes, of which only *HOXD13* is associated a cluster of reverse-oriented CBS elements (*CBS8-12*). By contrast, *HOXD8* and *HOXD9* have two forward-oriented CBS elements (*CBS13,14*) and *HOXD10-12* and *HOXD1-4* have no CBS element (Figs. 5a-c and Supplementary Fig. 6a). We performed Hi-C and found that there are strong long-distance chromatin interactions between the human *HOXD13* and the C-DOM left boundary, consistent with the convergent rule of forward-reverse CBS elements (Extended Data Fig. 10a,b). Interestingly, within the left boundary, there are two outward-oriented CBS elements (*CBS3* and *CBS5*, Fig. 5b and Supplementary Fig. 6b).

**Fig. 5.**
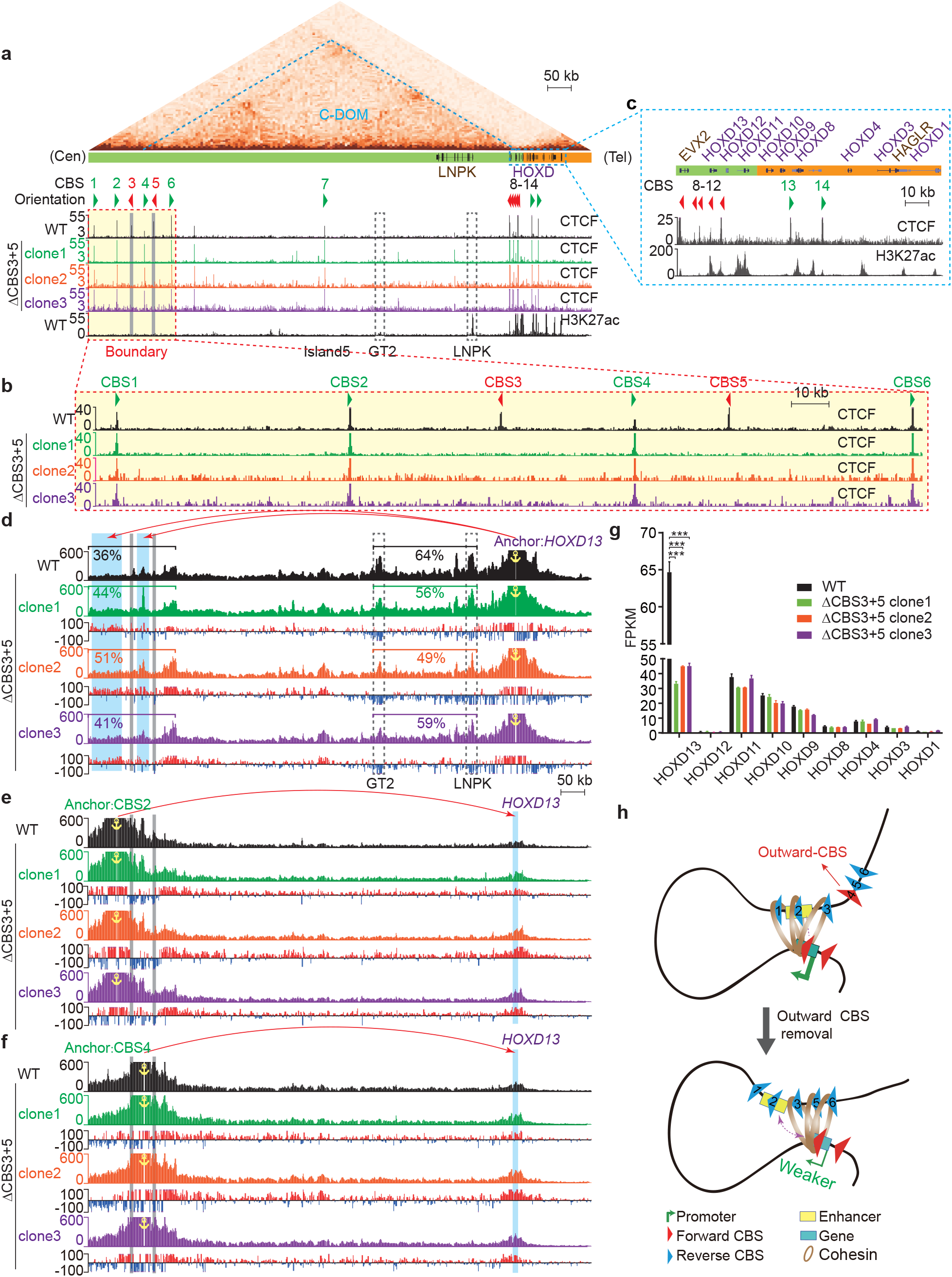
Double knockout of outward-oriented *CBS3* and *CBS5* downregulates *HOXD13*. **a**, CTCF and H3K27ac ChIP-seq profiles of the *HOXD* centromeric regulatory TAD (C-DOM) in wild-type (WT) and in *CBS3* and *CBS5* double knockout (Δ*CBS3*+*5*) homozygous single-cell clones. The left boundary (red-dotted rectangle) comprises a cluster of CTCF sites, *CBS1-6*, of which *CBS3* and *CBS5* are in the reverse orientation. Arrowheads indicate CBS elements with orientations. **b**, Close-up of CTCF ChIP-seq profiles at the C-DOM left boundary marked by red dotted rectangle in **a**. Note the abolishment of CTCF peaks at the locations of *CBS3* and *CBS5* in the three single-cell deletion clones. **c**, Close-up of CTCF and H3K27ac ChIP-seq profiles at the *HOXD* cluster marked by blue dotted rectangle in **a**. Note the five reverse-oriented CBS elements associated with *HOXD13* but not others. **d**, 4C profiles using *HOXD13* as an anchor in Δ*CBS3*+*5* single-cell clones compared to WT clones, showing increased chromatin interactions with the left boundary and decreased chromatin interactions with the H3K27ac-enriched *GT2 or LNPK* element upon double knockout. **e-f**, 4C profiles using *CBS2* (**e**) or *CBS4* (**f**) as an anchor confirming increased chromatin interactions with *HOXD13*. **g**, Expression levels of *HOXD* genes in three Δ*CBS3*+*5* single-cell clones compared to WT clones. Data as mean ± SD, \**p* < 0.05, \*\**p* < 0.01, \*\*\**p* < 0.001; one-tailed Student’s *t* test. **h**, A model illustrating the regulatory role of outward oriented CBS elements in intraTAD chromatin contacts and gene expression. TAD boundary normally contains clustered CBS elements with complex orientations of both divergence and convergence. The outward oriented CBS elements function as an important barrier to ensure productive intraTAD promoter-centered contacts. Loss of outward oriented CBS elements results in aberrant interTAD promoter-anchored chromatin interactions, leading to decreased gene expression.

We screened single-cell clones for deletion of both *CBS3* and *CBS5* elements and obtained three homozygous double-knockout cell clones (Δ*CBS3+5*, Supplementary Fig. 6e). We first confirmed the loss of CTCF enrichments at both *CBS3* and *CBS5* (Fig. 5a,b). We then performed 4C with *HOXD13* as an anchor and found that there is a significant increase of chromatin interactions with the left boundary upon deletions of the two outward-oriented CBS elements (Fig. 5d). This increased chromatin interactions were confirmed by reciprocal 4C experiments with *CBS2* or *CBS4* as an anchor (Fig. 5e,f). By contrast, there is a significant decrease of chromatin interactions with proximal sequences (Fig. 5d). Finally, we performed RNA-seq experiments and found that deletion of outward-oriented CBS elements results in a significant decrease of expression levels of *HOXD13* but not *HOXD12-1*, indicating that the two outward-oriented CBS elements play a key role in *HOXD13* gene regulation (Fig. 5g and Supplementary Fig. 6f).

## Discussion

The enormous connectivity of billions of neurons are intricately related to the precise cell-specific expression patterns of *cPcdh* genes in the brain. Specifically, the isoform-promiscuous *cis* hetero-dimerization and isoform-specific *trans* homo-dimerization of combinatorially expressed cPcdhs mediate cell-specific recognition between neurons, resulting in the ingenious dendrite self-avoidance and axonal coexistence^27,35-42^. Intriguingly, chromatin establishes a stable *cPcdh* expression pattern for single neurons in the brain during human fetal development^43^. In addition, the proper spatiotemporal expression of the *HoxD* genes is central for limb development^29,44^. Here, we first comprehensively mapped *cPcdh* expression patterns in the neocortex during mouse brain development. We then dissected, through sequential and combinatorial deletion of candidate enhancer elements, how *cPcdhs* are regulated by a downstream clustered CTCF boundary super-enhancer. In addition, we further deleted outward-oriented CBS elements within clustered CTCF TAD boundaries of the *cPcdh* and *HOXD* loci. In conjunction with chromosome conformation and gene expression analyses, we found that outward-oriented CBS elements are crucial for intraTAD promoter-enhancer interactions and proper gene regulation (Fig. 5h).

The characteristics of the clustered CTCF TAD boundaries and their exact nature of insulation activities are under intense investigation^13,18,20,22,29,44-48^. The CBS elements within a clustered CTCF TAD boundary between *Pax3* and its distal enhancer cooperate redundantly for robust insulation in a largely orientation-independent manner^18^. In addition, various combinations of clustered CBS elements in distinct orientations could function as functional insulators when inserted between distal enhancers and their target promoters^20,22,49^. Moreover, the CBS elements within a clustered CTCF TAD boundary located between the C-DOM and T-DOM (telomeric domain) of *HoxD* have various functions for controlling its spatiotemporal gene expression patterns during limb development^29,44^. Here we show that the outward-oriented CBS elements of the left boundary of C-DOM are crucial for maintaining *HOXD13* gene expression (Fig. 5). Finally, consistent with the crucial role of outward-oriented CBS elements, an inversion leading to increased numbers of outward-oriented CBS elements within a clustered CTCF TAD boundary results in enhanced insulation activity and altered gene expression^50^.

CTCF plays a key role in the formation of insulation at TAD boundaries, where in many cases multiple CBS elements are arranged in a clustered array with complex orientations, such as the telomeric boundary of the *Pcdhβγ* TAD and the centromeric boundary of the C-TOM at the *HOXD* cluster. According to the convergent rule, the inward-oriented CBS elements within the clustered CTCF TAD boundaries are essential for intraTAD chromatin interactions^8,9^. Thus, chromatin contacts such as long-distance enhancer-promoter interactions primarily occur within the TADs^51,52^. However, recent studies provide increasing evidence for interTAD communications between neighboring TADs^50^. Although TADs are likely a statistical model from population cells, there is clear evidence of clustering of CBS elements at TAD boundaries^13,15,48^. On one hand, the outward-oriented CBS elements could be crucial for intraTAD chromatin interactions. On the other hand, the outward-oriented CBS elements function as crucial insulators blocking aberrant chromatin interactions from neighboring TADs as we observed a sharp edge for increased outward chromatin interactions (Fig. 4).

The structure of TADs and loops is closely related to gene expression. Their formation is thought to be resulted from CTCF-anchored active cohesin “loop extrusion”. Specifically, cohesin extruded loops anchored between convergent forward-reverse CBS elements can enhance or stabilize enhancer-promoter spatial contacts, which may or may not activate gene expression^53-56^. In addition, both the spatial and linear distances between distal enhancers and target promoters could be crucial for gene regulation^57-59^. Our study on *Pcdhγc3* regulation is consistent with this idea. *Pcdhγc3*, the most abundantly expressed *cPcdh* isoform in the mouse neocortex (Extended Data Fig. 1e), plays a crucial role in dendrite arborization^34^. It is puzzling why *HS5-1bL* deletion leads to a significant increase of expression levels of only *γc3* but not other members of *cPcdh* genes (Figs. 1i,2e and Extended Data Fig. 3e). It has been shown that gene activation is not only dependent on promoter associated CBS elements, but also on the density of enhancers in the genomic context^32,57^. Consistently, only *γc3*, but not *γc4* and *γc5*, is associated with two forward-oriented CBS elements (Fig. 2a) despite the fact that, compared with other members of *Pcdhβγ* clusters, all of the three *Pcdhγc* genes are located close to the downstream distal enhancers of *HS18-20* which contain two reverse-oriented CBS elements (Fig. 1b). A “double clamp” interaction mechanism of the two forward-reverse CBS pairs, similar to the activation of members of the *Pcdhα* gene cluster, may be utilized to activate *γc3*. This explains why only *γc3*, but not *γc4* and *γc5*, expression levels are enhanced upon deletion of the *HS5-1bL* (*CBSbc*) insulator.

Our data highlight a significant role of the outward-oriented CBS elements in providing insulation for restraining aberrant interTAD interactions and maintain proper intraTAD promoter-enhancer interactions and gene expression. *HOXD13* showed strong interactions with two H3K27ac-marked regions of *LNPK* and *GT2*. We observed a significant decrease of chromatin interactions with *LNPK* and *GT2* upon removal of outward-oriented CBS elements (Fig. 5d), explaining the decreased levels of *HOXD13* gene expression. This suggests that the outward-oriented CBS boundary elements of the C-TOM may ensure proper interactions with *LNPK* and *GT2* for precise control of *HOXD13* expression patterns.

We observed that H3K9me3 marks correlate with the monoallelic *cPcdh* gene expression and that H3K9me2 marks correlate with both monoallelic and biallelic *cPcdh* gene expression^26^. The correlation between H3K9me3 enrichments and DNA methylation levels in both monoallelic and biallelic *cPcdh* genes may be coordinated by UHRF1^60^. Interestingly, almost every members of the *cPcdh* genes is marked by both repressive (H3K9me3 and H3K9me2) and active (H3K36me3) marks (Fig. 3), suggestive of a facultative heterochromatic state related to monoallelic *cPcdh* gene expression. Whether the overlap of H3K9me3 and H3K36me3 is resulted from allelic differences of the epigenome and/or admixtures of distinct subpopulation cells awaits further exploration.

## Acknowledgements

We are grateful for technical supports from Leyang Wang, Mo Zhang, and Jingwei Li. This work was supported by grants from the National Natural Science Foundation of China (91940303), the National Key R&D Program of China (2022YFC3400200) and the Science and Technology Commission of Shanghai Municipality (19JC1412500 and 21DZ2210200).

## Author contributions

Q.W. conceived the research. X.G. and K.H. performed experiments. H.H. and X.G analyzed data. Z.W. and W.X. supervised the mouse experiments. X.G, H.H. and Q.W. wrote the manuscript with inputs from all authors.

## Competing interests

The authors declare no competing interests

## Data availability

All raw high-throughput sequencing data and processed files could be accessed via: https://www.ncbi.nlm.nih.gov/geo/query/acc.cgi?&acc=GSE210817.

## Code availability

No code or model was generated in this study.

## Notes

### Competing Interest Statement

The authors have declared no competing interest.

