## Supplementary material for "Outward-oriented sites within clustered CTCF boundaries are key for intraTAD chromatin interactions and gene regulation": Methods and Supplementary Figure

### Cell culture

Human HEK293T cells were cultured in DMEM (Hyclone) supplemented with 10% (v/v) FBS (Gibco) and 10 U/ml penicillin-streptomycin (Gibco 15140122). Human Hec1B cells were cultured in MEM (Hyclone) supplemented with 200 mM L-glutamine, Earle's balanced salt solution (Hyclone SH30024.01), 1 mM sodium pyruvate (Sigma SLBP4879V), 10% fetal bovine serum, and 10 U/ml penicillin-streptomycin. All cells were incubated at 37 °C and 5% $CO_2$ in a humidified incubator, and passaged every three days.

### CRISPR screening of CBS-deletion single-cell clones

*CBS3* and *CBS5* within the clustered CTCF TAD boundary of the *HOXD* locus were sequentially knocked out by two steps of CRISPR/Cas9-mediated DNA fragment editing (Supplementary Table 2)[9,61]. In the first step, *CBS3* or *CBS5* was deleted in HEK293T cells to generate homozygous *CBS3* or *CBS5* knockout cell clones. In the second step, a *CBS3* knockout cell clone was used to delete *CBS5* and a *CBS3* knockout cell clone was used to delete *CBS3*, to generate cell clones homozygous for *CBS3* and *CBS5* double knockout (ΔCBS3+5).

For each step of CRISPR editing, cells were grown to ~80% confluency in 6-well tissue culture plates and transfected with 1.5 µg of pcDNA3.1-Cas9 and 1.5 µg dual sgRNA expression plasmids per well by Lipofectamine 3000 (Invitrogen, L3000015). Two days after transfection, 2 µg/ml puromycin was added to the cell growth medium and maintained for 5 days to select the transfected cells. The transfected cells were suspended into single cell solutions, diluted and plated into 96 well plates. Two weeks later, single-cell clones were picked and genotyped by PCR with specific primers (Supplementary Table 1). Positive clones were Sanger sequenced.

The sgRNA expression plasmids were constructed as previously described[9,61] with paired oligonucleotides (Supplementary Table 1). For each sgRNA, a forward oligonucleotide with the (ACCG-5') overhang and a reverse complementary oligonucleotide with the (AAAC-5') overhang were annealed and cloned into the BsaI site of pGL3-U6-sgRNA-PGK-Puro vector for sgRNA transcription[62]. All sgRNA-expressed plasmids were confirmed by Sanger sequencing.

### ***In vitro* synthesis of Cas9 mRNA and sgRNAs for micro-injection**

A T7 promoter-driven Cas9 expression plasmid[63] was linearized by XbaI, purified and *in-vitro* transcribed using the mMESSAGE mMACHINE T7 Ultra Kit (ThermoFisher, AM1345). The transcribed Cas9 mRNA was treated with Turbo DNase to remove template DNA and added with poly-A tail. The sgRNA template was generated by PCR amplification with specific primers (Supplementary Table 1) to contain a T7 promoter, a 20 nt target sequence, and a scaffold region[49]. The amplified product was purified and transcribed using the MEGAshortscript T7 Kit (ThermoFisher, AM1354). The transcribed sgRNA was treated with Turbo DNase. Both sgRNA and Cas9 mRNA were purified by the MEGAclear Kit (ThermoFisher, AM1908), eluted into elution buffer and stored at -80°C before usage.

### **Generation of CBS-deletion mice**

Mice were maintained in an SPF mouse facility at 23 °C with a 12 h (7:00-19:00) light /12 h (19:00-7:00) dark cycle. All the mouse experiments were approved by the Institutional Animal Care and Use Committee (IACUC) of Shanghai Jiao Tong University (Protocol#: 1602029).

C57BL/6J mice were used as embryo donors and ICR mice were used as foster mothers. 6-week-old C57BL/6J female mice were super-ovulated and mated with the sterilized C57BL/6J male mice to produce enough embryos.

Zygotes were collected from the oviducts of the super-ovulated C57BL/6J female mice, washed with 200 μg/ml hyaluronidase (Sigma, V900833) and incubated in the M2 medium (Sigma, M7167) in a 5% $CO_2$ incubator at 37 °C for 1 h. Viable embryos were then microinjected with a solution containing 100 ng/μl of Cas9 mRNA and 50 ng/μl each of two sgRNAs targeting each fragment and recovered in a 5% $CO_2$ incubator at 37 °C for 1 h. The injected live embryos were transplanted into the oviducts of the pseudo-pregnant ICR female mice under a stereoscopic microscope, with each ICR mouse receiving 25-30 embryos. Post-surgery mice were maintained at a 37 °C heating plate for 1 h for recovery before being transferred back for regular housing until the birth of the F0 mice. Mice exhibiting signs of dystocia at day 20 received C-section rescue.

The chimeric F0 mice were screened ten days later for desired deletions by PCR genotyping with specific primers (Supplementary Table 1) and the chimeric mutant mice were confirmed by Sanger sequencing. The F0 mice of 2-month old with desired deletions were mated with wildtype C57BL/6J mice to generate heterozygous F1 mice. The F1 mice were genotyped by PCR. The targeted F1 male and female mice were crossed to generate the F2 homozygous mice, which were used for downstream experiments. The wildtype F2 littermates were used as controls.

**ATAC-seq**

ATAC-seq was performed following the omni-ATAC procedure[64] with some modifications. Briefly, the neocortex was microdissected and digested with 0.00625% collagenase followed by filtration through a 100 μm cell strainer to generate single-cell suspensions. The suspended cells were washed twice with PBS solution and then counted. $10^5$ cells were aliquoted and resuspended with 50 μl of the resuspension buffer (RSB) supplemented with 0.1% NP40, 0.1% Tween-20, and 0.01% digitonin followed by incubation on ice for 3 minutes for lysis. The nuclei were then washed with 1 ml of RSB supplemented with only 0.1% Tween-20 and resuspended in 50 µl of transposition mix, which includes 25 µl of 2× TD buffer, 3 µl transposase, 16 µl PBS, 0.5 µl 1% digitonin, 0.5 µl 10% Tween-20 and 5 µl $H_2O$, followed by incubation at 37 °C for 30 min in a shaker at 1000 rpm. Transposition products were immediately cleaned up with 1× Ampure XP Beads (Beckman, A63881) and the purified DNA was used for library construction by the TruePrep DNA Library Prep Kit V2 for Illumina (Vazyme Biotech, TD502). The ATAC-seq libraries were pooled and sequenced at the 2x150 bp mode using an Illumina NovaSeq 6000 platform.

**RNA-seq**

Total RNA was extracted from cultured cells or the microdissected mouse neocortex using the Trizol reagent (ThermoFisher, 15596026). For each sample, 1 µg of total RNA was used to enrich mRNA using Poly(A) Magnetic Isolation Module (NEB, E7600). The mRNA was purified twice, eluted into the fragmentation buffer with random primers for fragmentation at 94 °C for 15 min, and reverse-transcribed into the first strand cDNA. The second strand was then synthesized by the second-strand DNA polymerase. The generated double-stranded cDNA was purified with Ampure XP beads (Beckman, A63881), end repaired, 3'-adenylated, and 5'-phosphorylated. The Illumina sequencing adaptors were then ligated to both ends of cDNA and excess adaptors were removed using Ampure XP beads (Beckman). The purified cDNA was amplified by PCR with barcoding to generate RNA-seq libraries. Pooled multiplexed libraries were sequenced at the 2x 150 bp mode on an Illumina NovaSeq 6000 platform. All experiments were performed with at least two biological replicates.

Fetal sex is determined based on the expression of *Xist* and *Ddx3y*. For expression mapping at different embryonic states during cortical development, mouse neocortices were isolated from both males (*Xist*$^+$, *Ddx3y*$^-$) and females (*Xist*$^-$, *Ddx3y*$^+$). As we did not observe sex-specific differences in *cPcdh* expression at any developmental stages (Extended Data Fig. 1f,g), we did not discriminate between male and female mouse neocortices for all the following experiments.

**ChIP-seq**

Mouse neocortical tissues was microdissected and dissociated into single cells using 0.00625% collagenase. 2x $10^6$ of mouse neocortical or cultured human cells were aliquoted and crosslinked using 1% formaldehyde in PBS containing 10% FBS for 10 min at room temperature followed by two washes with PBS containing proteinase inhibitors (Roche, 04693132001). Cells were then resuspended and lysed twice with 1 ml of prechilled lysis buffer containing proteinase inhibitors (Roche) for 10 min with slow rotations. For cortical cells, additional SDS was added to the lysis solution to adjust the final SDS concentration to 0.4% for better sonication. Cells were then sonicated at 25% power for a train of 15 s ON and 30 s OFF for 20 cycles with the Bioruptor system to break the genomic DNA into 0.1-10 kb fragments. After sonication, the SDS-free lysis buffer was added to cortical samples to dilute the SDS concentration to 0.1% to allow the binding of antibodies to proteins. After removal of the insoluble debris by centrifuging at 14,000 g for 10 min, the lysate was pre-cleaned with protein-A agarose beads (Millipore, 16-157) for 1 h with slow rotations to remove non-specific binding and then incubated with specific antibody (Supplementary Table 3) overnight with slow rotations at 4°C. The antibody-protein-DNA complex was isolated by adding protein A-agarose beads and incubated for 4 h with slow rotations followed by sequential washes with the low-salt buffer, high-salt buffer, salt-free buffer, LiCl buffer, and TE buffer. The beads were eluted in the elution buffer (0.1 M $NaHCO_3$, 1% SDS) and de-crosslinked by proteinase K at 65 °C overnight. The DNA was finally purified and used for the construction of DNA library using the VAHTS Universal DNA

Library Prep Kit for Illumina V3 (Vazyme Biotech, ND607). ChIP-seq libraries were sequenced at the 2x 150 bp mode on an Illumina NovaSeq 6000 platform.

**4C**

4C experiments were performed as previously described[65]. In brief, mouse neocortices were microdissected and dissociated with 0.00625% collagenase to obtain single cells. Mouse neocortical or cultured human cells were crosslinked in 2% formaldehyde for 10 min at room temperature and quenched by adding excess pre-chilled glycine. The fixed cells were incubated on ice for 5 min, and permeabilized twice with the permeabilization buffer (50 mM Tris–HCl pH 7.5, 150 mM NaCl, 5 mM EDTA, 0.5% NP-40, 1% Triton X-100, and protease inhibitors), with slow rotations for 10 min at 4 °C. The permeabilized cells were then digested with DpnII for 16 h at 37 °C while shaking at 900 rpm followed by incubation at 65 °C for 20 min to inactive DpnII. Proximity ligation was then performed for 16 h at 16°C by adding T4 DNA ligase in 1x T4 ligation buffer. The ligated product was de-crosslinked by proteinase K at 65 °C overnight, treated with RNase A at 37 °C for 45 min, and purified with phenol-chloroform followed by ethanol precipitation. DNA was sonicated to 0.2-1 kb fragments using the Bioruptor system at 33% power for a train of 15 s ON and 30 s OFF for 4 cycles.

The anchor fragments were linear-amplified using a specific 5' biotin-tagged primer (Supplementary Table 1) with 120 cycles (95 °C 20 s for denaturing, 60 °C 20 s for annealing, and 72 °C 90 s for elongation). The amplified products were finally denatured at 95 °C for 5 min and then immediately chilled on ice to prevent reannealing. The biotin-tagged single-stranded DNA (ssDNA) was pulled down with Streptavidin Magnetic beads (ThermoFisher, 65001), and ligated with adaptors which were generated by annealing two partially complementary single-stranded oligonucleotides (Adapter-U and Adapter-L, Supplementary Table 1) by T4 DNA ligase in 1x T4 ligation buffer containing PEG8000 at 16 °C for 12 h. The on-bead ligation products were washed twice with the B/W buffer (5 mM Tris-HCl pH 7.5, 1 M NaCl, and 0.5 mM EDTA) to remove excess adaptors and used as the template for PCR amplification of the 4C libraries with barcoded anchor-specific P5-forward primers and indexed P7-reverse primers (Supplementary Table 1). Multiplexed libraries from the same anchor with different combinations of barcodes and indexes were pooled for purification with a PCR purification kit (Qiagen, 11732668001). 4C libraries were sequenced at the 2x 150 bp mode on an Illumina NovaSeq 6000 platform.

***In situ* Hi-C**

*In situ* Hi-C experiments were performed as previously described[8]. Briefly, 5x $10^6$ cells were cross-linked with formaldehyde and incubated in a lysis buffer (10 mM Tris-HCl pH8.0, 10 mM NaCl, 0.2% NP-40) to obtain nuclei. Nuclei were permeabilized with SDS, quenched by adding Triton X-100, and digested with MboI (NEB, R0147M) overnight at 37 °C, followed by heat inactivation at 62 °C. The ends of restriction fragments were filled in with biotin-14-dATP (Thermo, 19524016), dCTP, dTTP, and dGTP using the Klenow fragment of DNA polymerase I (NEB, M0210L), and ligated using T4 DNA ligase (NEB, M0202S). After ligation, the nuclei were de-crosslinked overnight at 68 °C. The ligated DNA was precipitated with ethanol and fragmented by sonication. 300-500 bp DNA fragments were selected using AMPure XP beads (Beckman, A63881). The biotinylated DNA was pulled down with M-280 Streptavidin beads (Thermo, 11206D), washed, and used for library construction on bead. The DNA was end-repaired, tailed with dA using Klenow exo$^-$ (NEB, M0212S), and ligated with Illumina U-type adaptors, followed by adaptor linearization with a USER enzyme (NEB, M5505S). The obtained DNA was used as templates for PCR amplification with 10-12 cycles. PCR products were purified with AMPure XP beads (Beckman, A63881) and sequenced at the 2x 150 bp mode on an Illumina NovaSeq 6000 platform.

### High-throughput sequencing data analysis

RNA-seq reads were aligned against the mouse (GRCm38) or human (GRCh38) genome using Hisat2[66] (version 2.0.4) to generate the sequence alignment map (SAM) files. SAM files were then converted into the binary versions (BAM files) using Samtools (version 1.15.1). BAM files were used as input for Cufflinks[67] (version 2.1.1) to calculate FPKM values.

ChIP-seq and ATAC-seq reads were aligned to the mouse (mm9) or human (hg19) genome using Bowtie2 to generate SAM files, which were converted into BAM files using Samtools (version 1.15.1). BAM files were indexed and converted into BedGraph files.

For 4C data, P7 reads were initially demultiplexed based on the unique barcode-index combinations. The anchor primer sequences were trimmed and PCR duplicate reads were removed using FastUniq (version 1.1) of the UNIX system. The obtained unique reads were aligned against the mouse (mm9) or human (hg19) genome using Bowtie2 to generate SAM files, which were converted into BAM files. The BAM files were used as input for r3Cseq (version 1.20) in the R package (3.3.3) to calculate the reads per million (RPM) values.

Hi-C reads were pre-processed with HiC-Pro[68], including reads alignment to the human hg19 reference genome, interaction matrix generation, and ICE normalization. 10 kb ICE normalized interaction matrix was generated. TADs were called at a 10-kb resolution using the directionality index (DI) method[51]. Insulation scores were calculated at a 10-kb resolution as previously described[69].

Extended Data Fig1

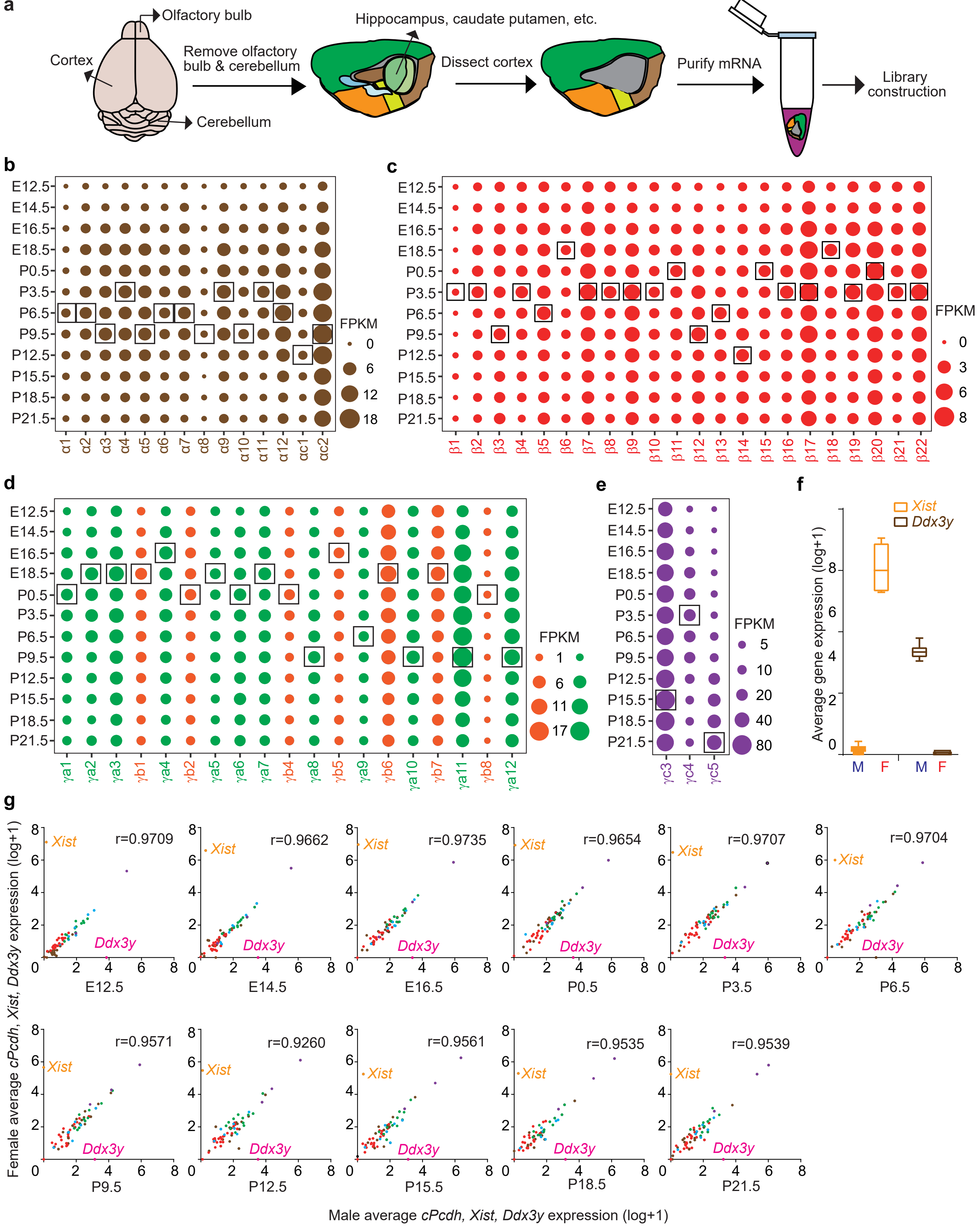

**Extended Data Fig. 1 | Expression patterns of the *cPcdh* genes during mouse neocortical development.** **a**, Schematics of the procedure for generating RNA-seq libraries of the mouse neocortex. Mouse neocortex was microdissected from the whole brain by removing olfactory bulb and other tissues and lysed in Trizol reagents for mRNA purification and library construction. **b**-**e**, Expression patterns of members of the *Pcdh*$\alpha$ (**b**), *Pcdh*$\beta$ (**c**), and *Pcdh*$\gamma$ (**d**,**e**) clusters during mouse neocortical development. Dot size indicates the expression levels. The outlined dots indicate the peak expression of each *cPcdh* member at a specific time point. Expression levels were measured by RNA-seq. **f**, Normalized expression levels of gender markers of *Ddx3y* and *Xist* between male and female neocortices at P0.5. **g**, No gender bias in expression patterns of the *cPcdh* genes. Correlation between male and female *cPcdh* expression levels at all analyzed developmental stages in the mouse neocortex. Pearson correlation coefficients are indicated on the right corner. The Y-chromosome gene *Ddx3y* and X-chromosome gene *Xist* are shown to indicate male and female, respectively.

Extended Data Fig2

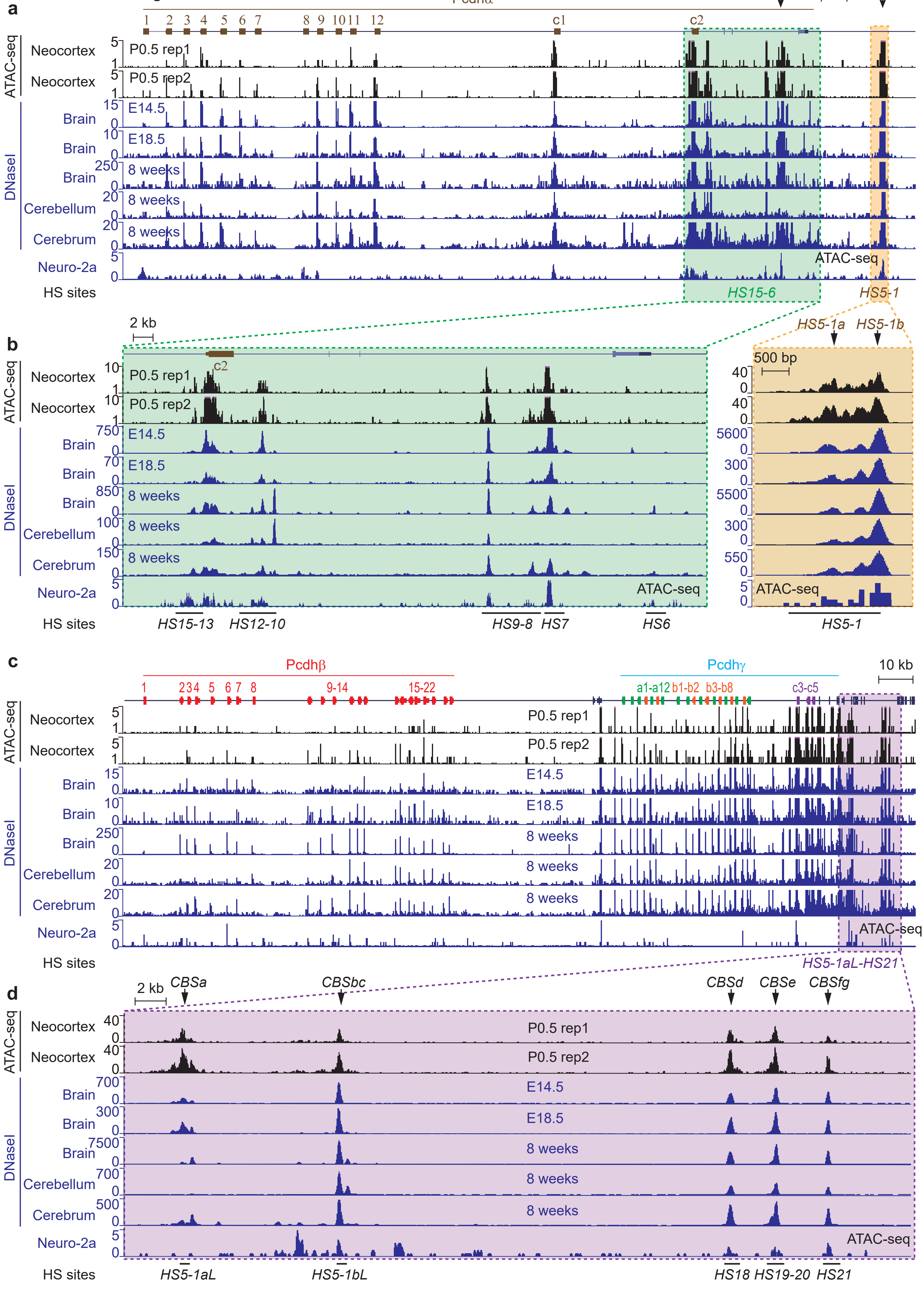

**Extended Data Fig. 2 | ATAC-seq profiles of the three *Pcdh* clusters in mouse newborn neocortex compared to DNaseI-seq profiles in mouse neural tissues.** **a**, ATAC-seq profiles of the *Pcdh*$\alpha$ cluster in the P0.5 mouse neocortex compared to publicly available DNaseI-seq profiles in the whole brains of E14.5, E18.5, and 8-week-old mice, the cerebellar and cerebral tissues of 8-week-old mice, and ATAC-seq profiles in the Neuro-2a cell line. **b**, Close-up of ATAC-seq and DNaseI-seq profiles at *HS1-15* sites highlighted in **a**. **c**, ATAC-seq profiles of the *Pcdh*$\beta\gamma$ clusters in the P0.5 mouse neocortex compared to DNaseI-seq profiles in the whole brains of E14.5, E18.5, and 8-week-old mice, the cerebellar and cerebral tissues of 8-week-old mice, and ATAC-seq profiles in the Neuro-2a cell line. **d**, Close-up of ATAC-seq and DNaseI-seq profiles at *HS5-1aL*, *HS5-1bL,* and *HS18-21* highlighted in **c**. CBS elements were indicated above.

Extended Data Fig3

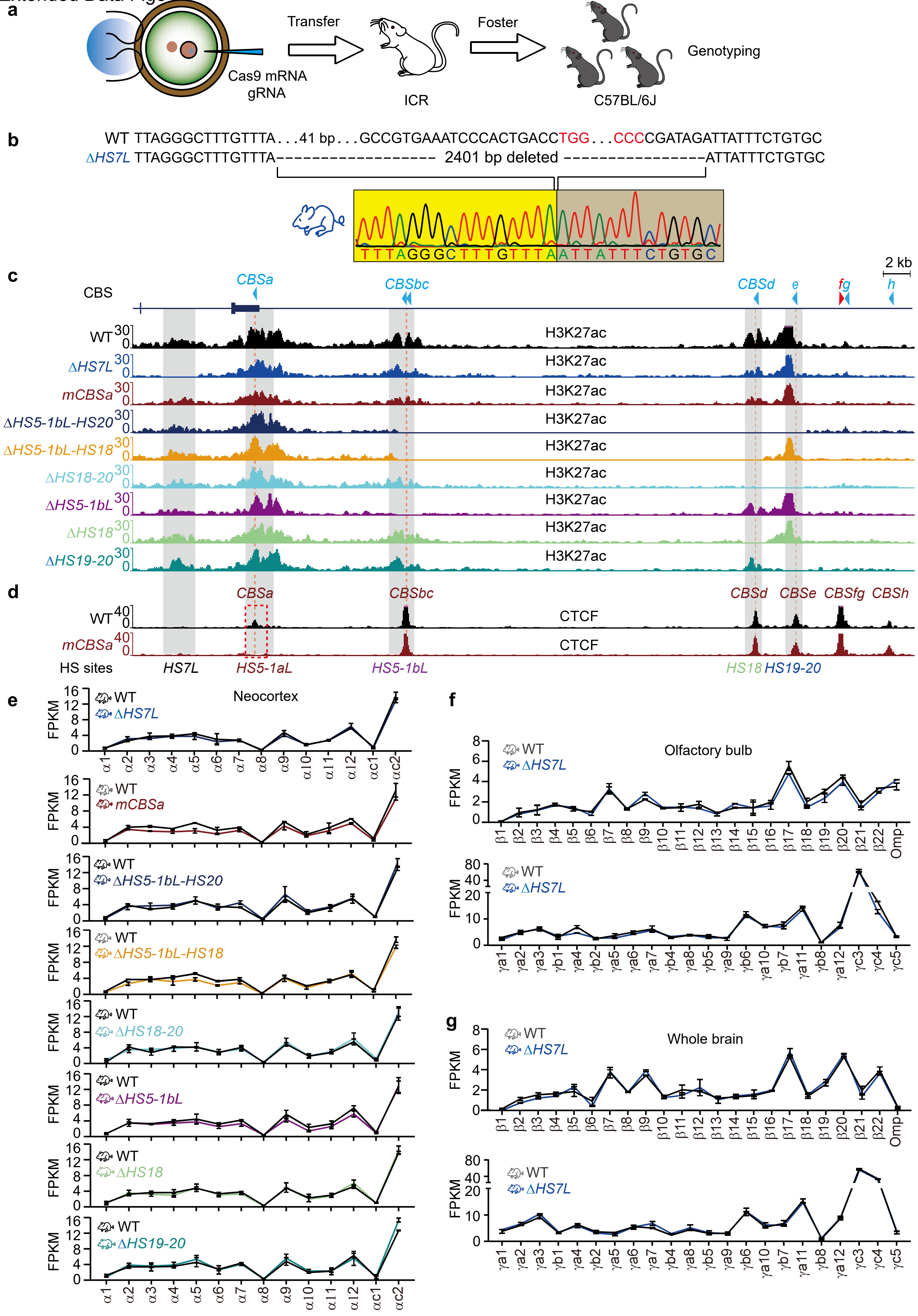

**Extended Data Fig. 3 | Mouse genetics of a repertoire of *cPcdh cis*-regulatory elements.** **a**, Schematics of the procedure for generating CRISPR/Cas9-based DNA fragment targeting mice. Zygotes from E0.5 C57BL/6J mice were microinjected with a solution containing Cas9 mRNA and sgRNAs targeting each fragment and transplanted into the oviducts of the pseudo-pregnant ICR female mice. The produced chimeric F0 mice were screened for desired deletions and crossed with wildtype C57BL/6J mice to generate heterozygous F1 mice. F1 mice were genotyped and crossed to generate F2 homozygous mice. **b**, Genotyping of *HS7L*-deleted (Δ*HS7L*) homozygous mice by Sanger sequencing. **c**, H3K27ac ChIP-seq profiles of the clustered CTCF TAD boundary in each mutant mouse line. **d**, ChIP-seq profiles showing the loss of CTCF enrichments at the *CBSa* element in *CBSa*-mutant (*mCBSa*) mice. **e**, RNA-seq showing no significant alteration of *Pcdh*$\alpha$ expression in all targeted mice. **f-g**, RNA-seq showing no alteration of *Pcdh* $\beta$ and $\gamma$ expression in the olfactory bulb and whole brain tissues of Δ*HS7L* mice. Omp, olfactory marker protein.

Extended Data Fig4

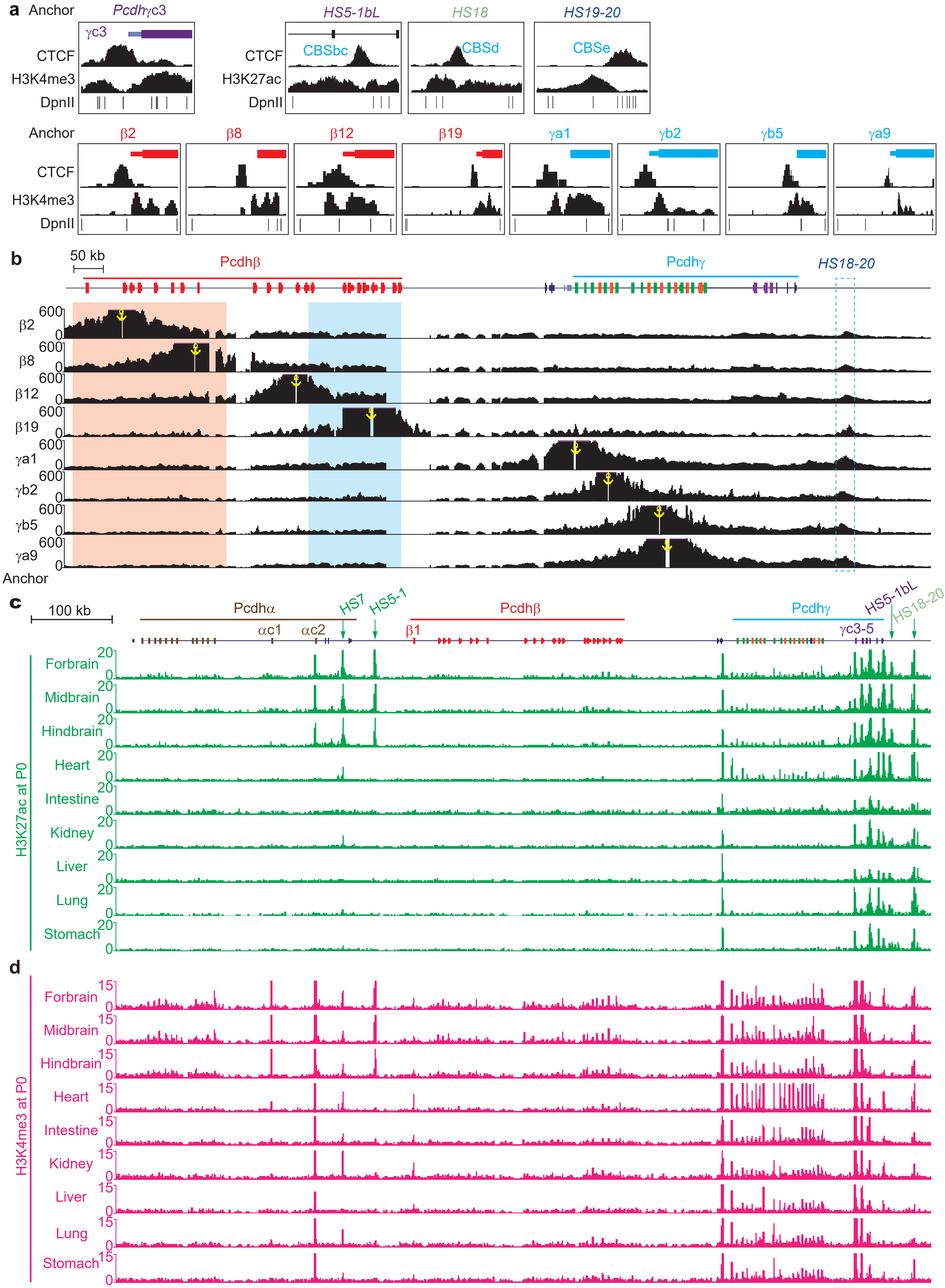

**Extended Data Fig. 4 | Chromosome conformation and tissue specificity of the *Pcdhβγ HS18-20* enhancers.** **a**, Schematic diagrams showing the positions of each 4C anchor. **b**, 4C profiles using a repertoire of *Pcdhβγ* promoters as anchors showing their close contacts with *HS18-20*. **c**,**d**, H3K27ac (**c**) and H3K4me3 (**d**) ChIP-seq profiles of histone marks in neural and non-neural tissues in P0 mice.

Extended Data Fig5

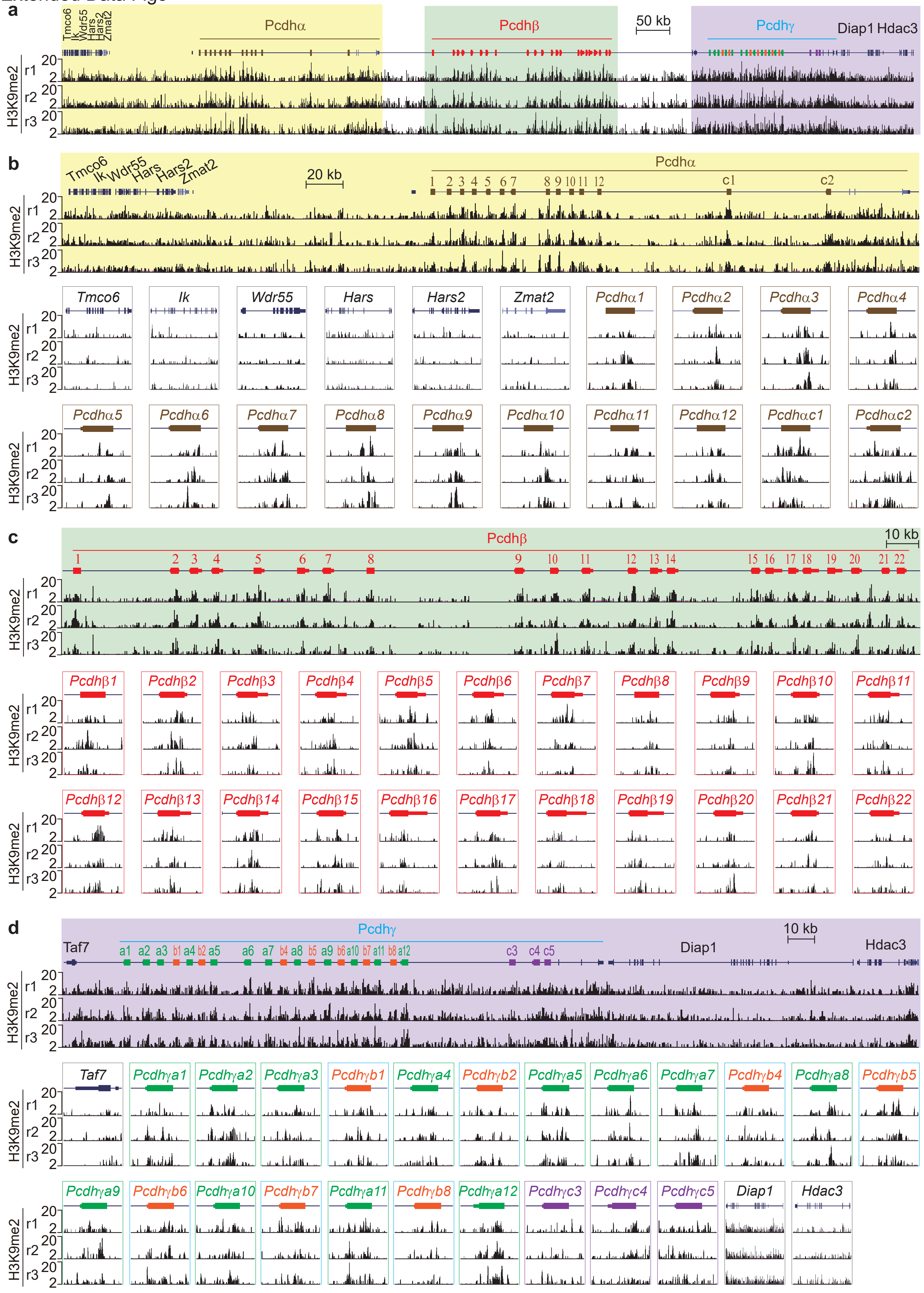

**Extended Data Fig. 5 | Enrichments of the heterochromatin mark of H3K9me2 at the *cPcdh* genes in the brain. a**, H3K9me2 ChIP-seq profiles at the *cPcdh* locus and its flanking regions with three replicates. **b-d**, Close-up of H3K9me2 profiles of the *Pcdh*$\alpha$ cluster (**b**), the *Pcdh*$\beta$ cluster (**c**), and the *Pcdh*$\gamma$ cluster (**d**) showing correlation of each c*Pcdh* variable exon with H3K9me2 compared to the flanking genes.

Extended Data Fig6

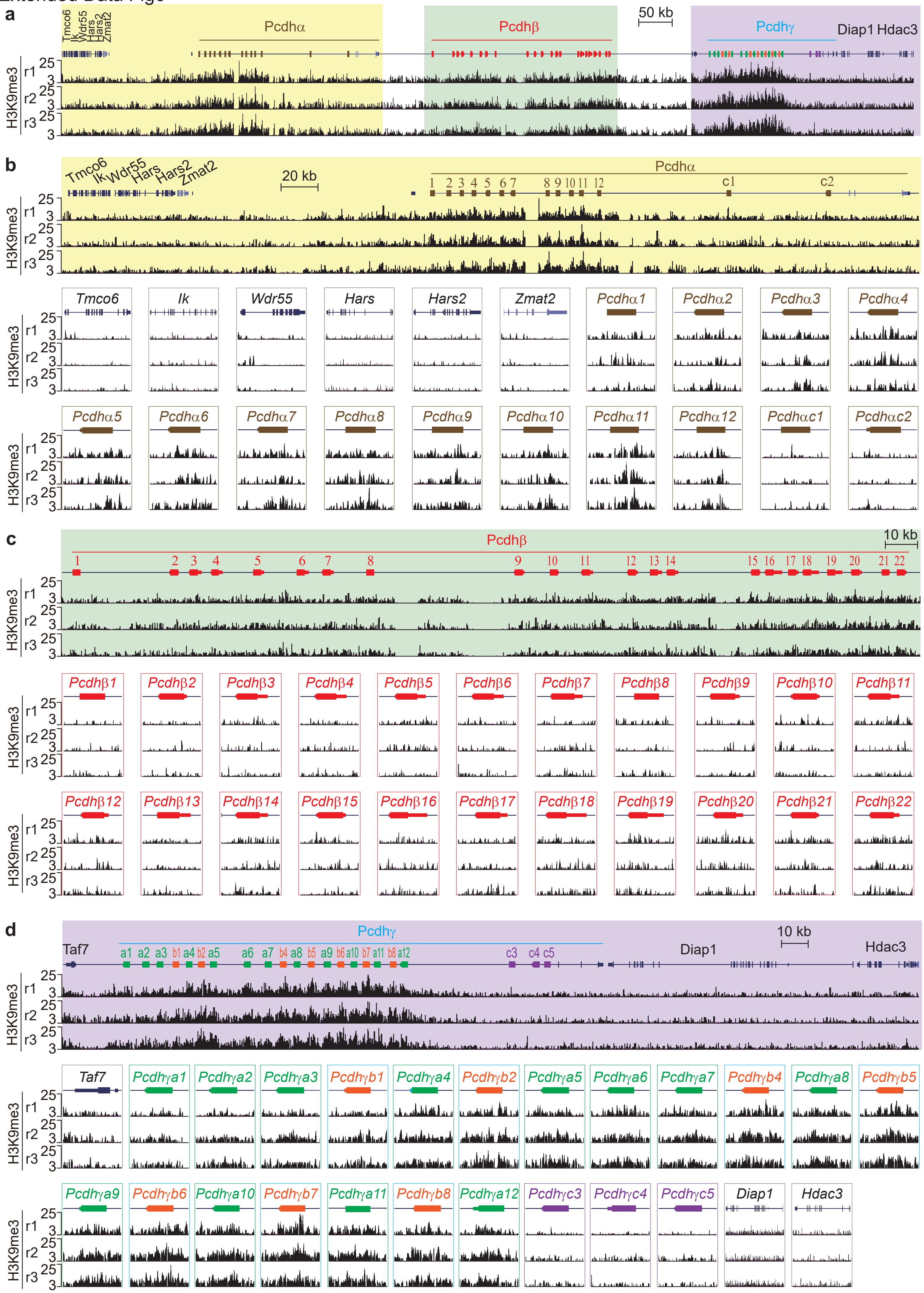

**Extended Data Fig. 6 | Enrichments of the heterochromatin mark of H3K9me3 at the monoallelic *cPcdh* genes in the brain. a**, H3K9me3 ChIP-seq profiles at the *Pcdh* locus and the flanking regions with three replicates. **b-d**, Close-up of H3K9me3 profiles of the *Pcdh*$\alpha$ cluster (**b**), within the *Pcdh*$\beta$ cluster (**c**), and downstream and within the *Pcdh*$\gamma$ cluster (**d**) showing correlation of each monoallelic *Pcdh* variable exon with H3K9me3. Note the absence of H3K9me3 mark in the five biallelic C-type *Pcdh* exons.

Extended Data Fig7

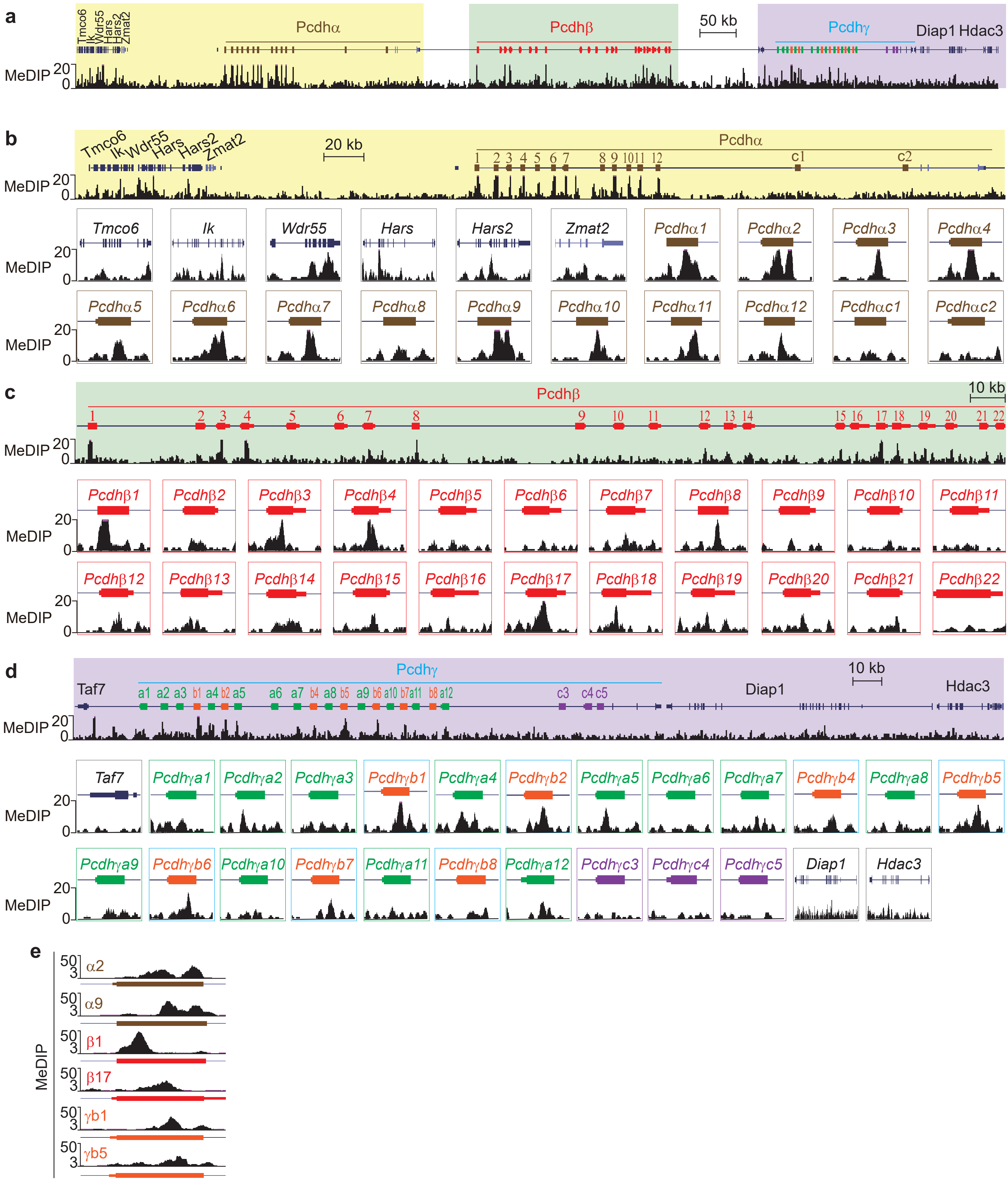

**Extended Data Fig. 7 | Enrichments of DNA methylation at the *cPcdh* genes in the brain.** **a**, MeDIP-seq profile at the *Pcdh* clusters and their flanking regions. **b-d**, Close-up of MeDIP-seq profiles of the *Pcdh*$\alpha$ cluster (**b**), the *Pcdh*$\beta$ cluster (**c**), and the *Pcdh*$\gamma$ cluster (**d**), showing the strong DNA methylation at the 3’ end of monoallelic *Pcdh*$\alpha$ variable exons and moderate DNA methylation at most *Pcdh*$\beta\gamma$ variable exons. **e**, Note the methylation at the 5’ end of *Pcdh*$\beta 1$ compared to methylation at the 3’ end of other *cPcdh* genes.

Extended Data Fig8

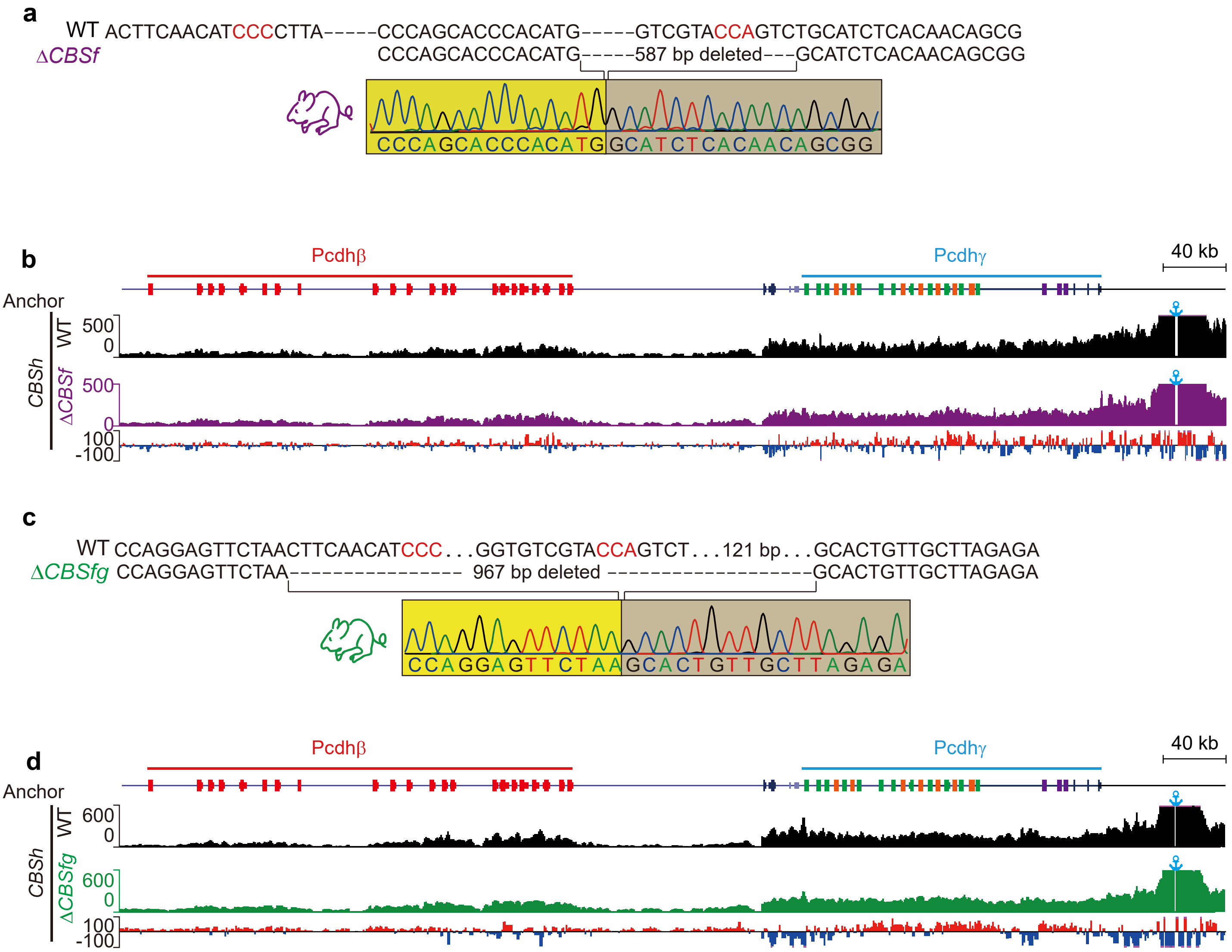

**Extended Data Fig. 8 | Increased chromatin interactions between *Pcdhβγ* and the downstream *CBSh* element upon the deletion of *CBSf* or *CBSfg*.** **a**, Genotyping of *CBSf*-deleted (Δ*CBSf*) homozygous mice by Sanger sequencing. **b**, 4C profiles using *CBSh* as an anchor, showing increased chromatin interactions with *Pcdhβγ* upon *CBSf* knockout. **c**, Genotyping of *CBSfg*-deleted (Δ*CBSfg*) homozygous mice by Sanger sequencing. **d**, 4C profiles using *CBSh* as an anchor, showing increased chromatin interactions with *Pcdhβγ* upon *CBSfg* knockout.

Extended Data Fig9

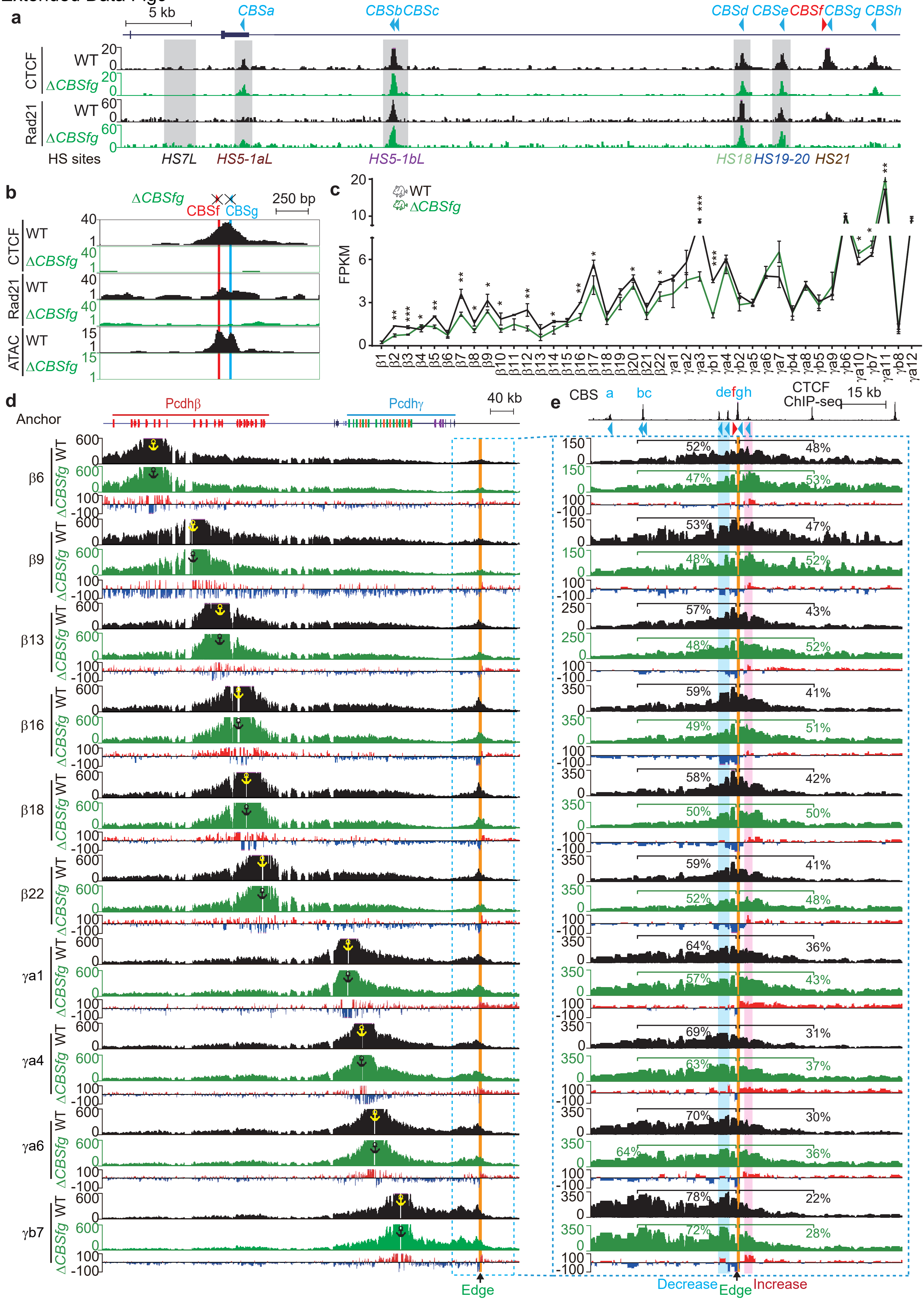

**Extended Data Fig. 9 | Combined deletion of both *CBSf* and *CBSg* elements results in a phenotype similar to that of the *CBSf* deletion. a**, CTCF and Rad21 ChIP-seq profiles of the downstream boundary of the *Pcdhβγ* TAD in *CBSfg*-deleted (Δ*CBSfg*) mice compared to their wild-type (WT) littermates showing the abolishment of CTCF and Rad21 binding upon *CBSfg* knockout. **b,** Close-up of CTCF and Rad21 ChIP-seq as well as ATAC-seq profiles at *CBSfg* in Δ*CBSfg* mice compared to their WT littermates. **c,** RNA-seq of *Pcdhβγ* in Δ*CBSfg* mice compared to their WT littermates showing a significant decrease of the expression of the *Pcdhβγ* clusters. Data as mean ± SD, $*p < 0.05$, $**p < 0.01$, $***p < 0.001$; one-tailed Student's *t* test. **d-e,** 4C profiles using a repertoire of *Pcdhβγ* promoters as anchors showing increased chromatin interactions beyond the location of *CBSfg* (highlighted in pink, **d** and **e**) and decreased chromatin interactions with *HS18-20* enhancers (highlighted in blue, **e**). Interaction differences (Δ*CBSfg* versus WT) are shown under the 4C profiles. Note a sharp transition edge at the location of the *CBSfg* element.

Extended Data Fig10

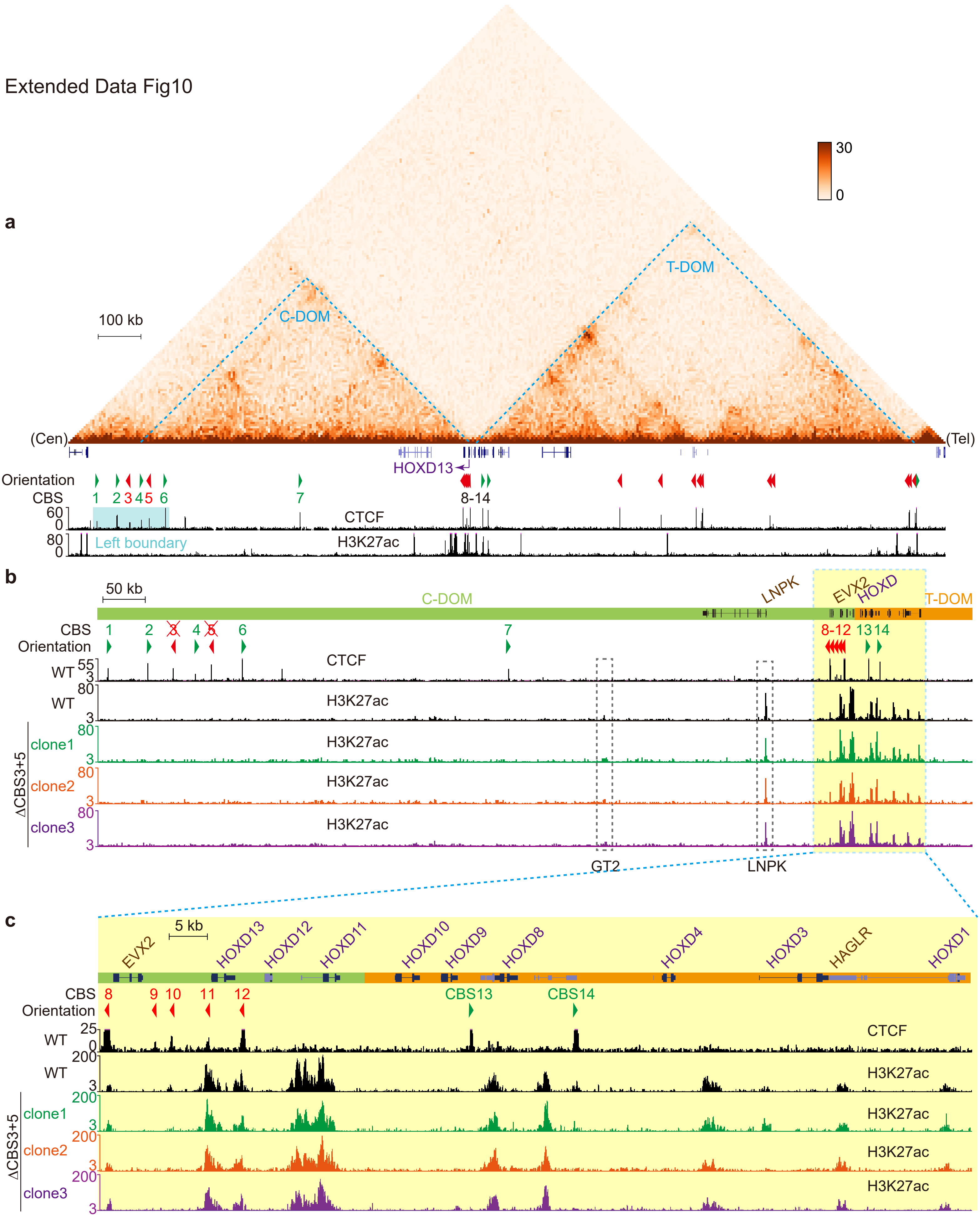

**Extended Data Fig. 10 | Outward-oriented *CBS* elements within the C-TOM boundary are crucial for *HOXD13* expression. a**, Hi-C map of the *HOXD* genomic region showing the TAD organization around the *HOXD* locus. The *HOXD* cluster are located between the two regulatory domains: centromeric TAD (C-DOM) and telomeric TAD (T-DOM). Arrowheads indicate CBS elements with orientations. CTCF and H3K27ac ChIP-seq profiles showing that the left boundary (highlighted in blue) of C-DOM is free of H3K27ac marks and comprises six clustered CBS elements (*CBS1-6*), of which *CBS3* and *CBS5* are outward-oriented. **b**, H3K27ac ChIP-seq profiles of the C-DOM in the three CBS*3* and *CBS5* double knockout (ΔCBS*3+5*) homozygous single-cell clones compared to wild-type (WT) clones. **c**, Close-up of the *HOXD* cluster outlined in **b** showing that *HOXD13* is associated with a cluster of five tandem reverse-oriented *CBS* elements.

Supplementary Fig1

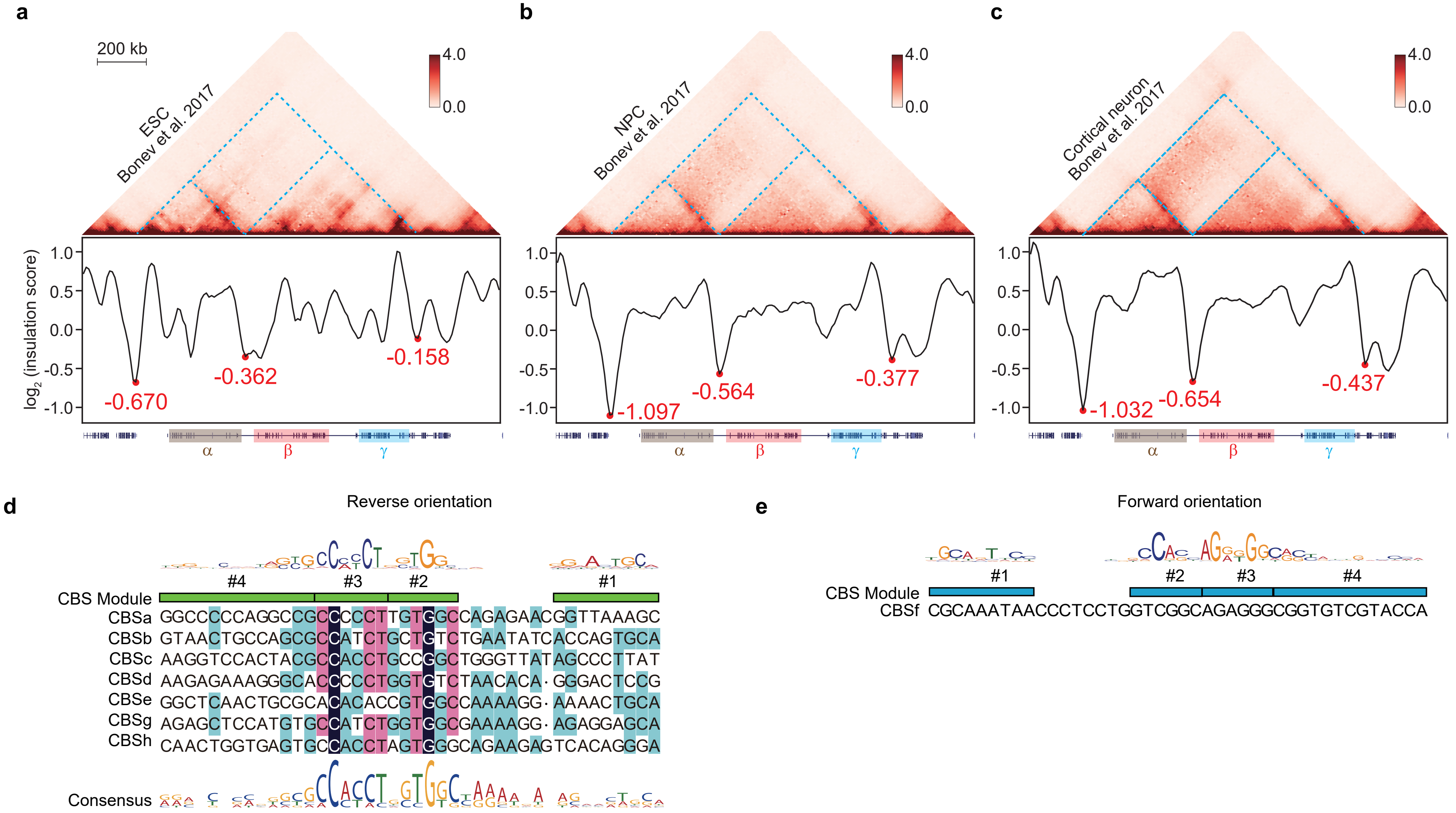

**Supplementary Fig. 1 | Organization of the three *cPcdh* clusters into a large megabase-sized superTAD during mouse neocortical development.** **a-c**, Hi-C maps of the three *cPcdh* clusters and its flanking regions in mouse ESCs (**a**), NPCs (**b**), and cortical neurons (**c**) obtained during *in vitro* differentiation of ES cells[70], as well as in mouse brain-derived cortical neurons[30] showing that the *cPcdh* superTAD containing *Pcdh*$\alpha$ and *Pcdh*$\beta\gamma$ TADs is already established in the mouse ES cells and reinforced during neural development. Insulation score values were shown under the HiC maps. Red dots represent the local minima of the insulation score at the *Pcdh*$\alpha$ and *Pcdh*$\beta\gamma$ TAD boundaries. **d**, An alignment of all seven reverse-oriented CBS elements (*CBSa-e, CBSg-h*) of the downstream boundary of *Pcdh*$\beta\gamma$ TAD downstream and sequences of the single forward-oriented *CBSf*. **e**, Sequences of the single forward-oriented *CBSf* of the downstream boundary of *Pcdh*$\beta\gamma$ TAD.

Supplementary Fig2

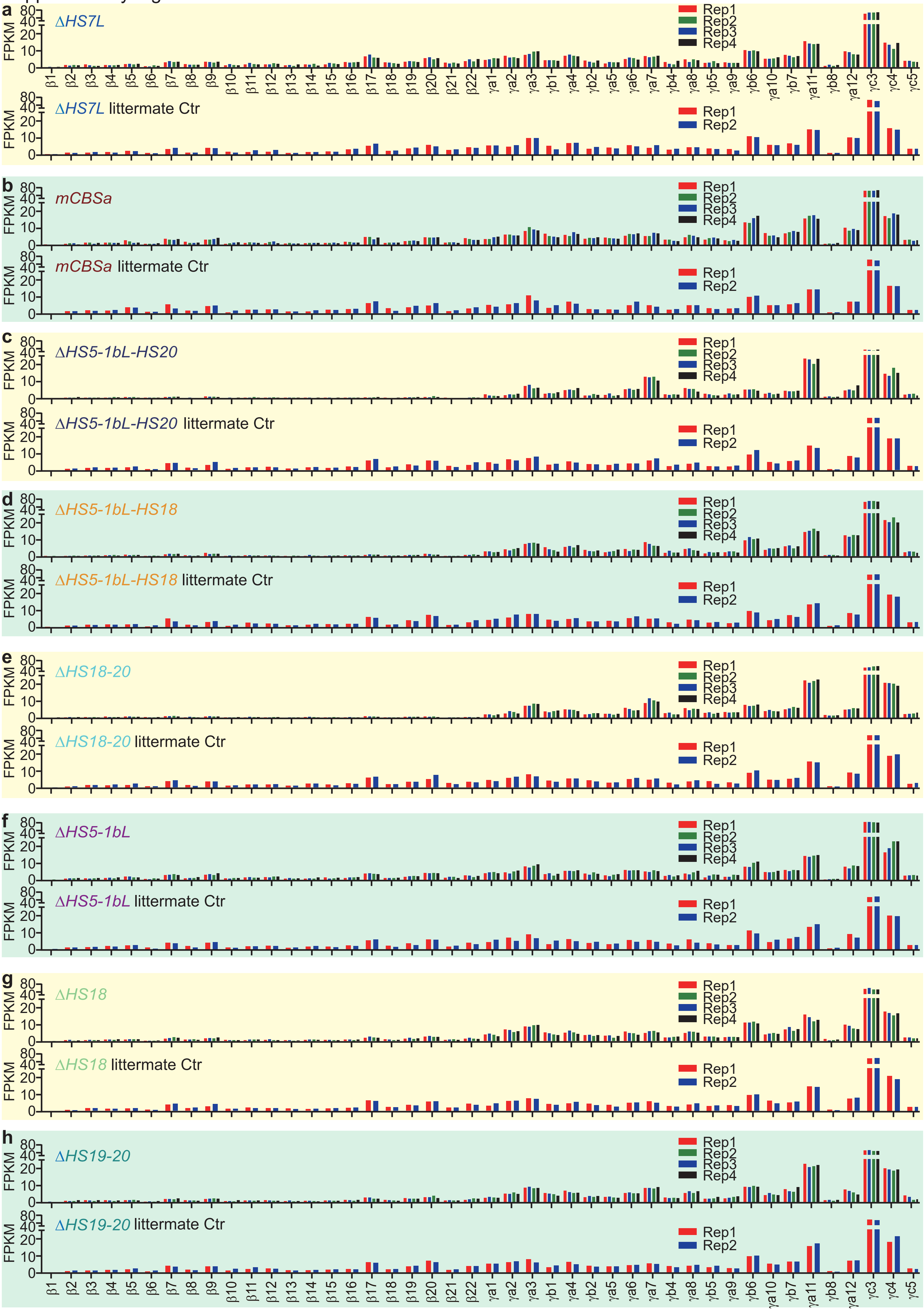

**Supplementary Fig. 2 | Replicates of RNA-seq levels of the *Pcdhβγ* genes.** **a-h,** Expression levels of replicates of RNA-seq experiments for Δ*HS7L* (**a**), m*CBSa* (**b**), Δ*HS5-1bL-HS20* (**c**), Δ*HS5-1bL-HS18* (**d**), Δ*HS18-20* (**e**), Δ*HS5-1bL* (**f**), Δ*HS18* (**g**), or Δ*HS19-20* (**h**) homozygous mice compared to their wild-type littermates. Expression levels were based on the FPKM values. Data as mean ± SD. * $P < 0.05$, ** $P < 0.01$, *** $P < 0.001$. For each mouse line, two wild-type replicates and four mutant replicates were performed.

Supplementary Fig3

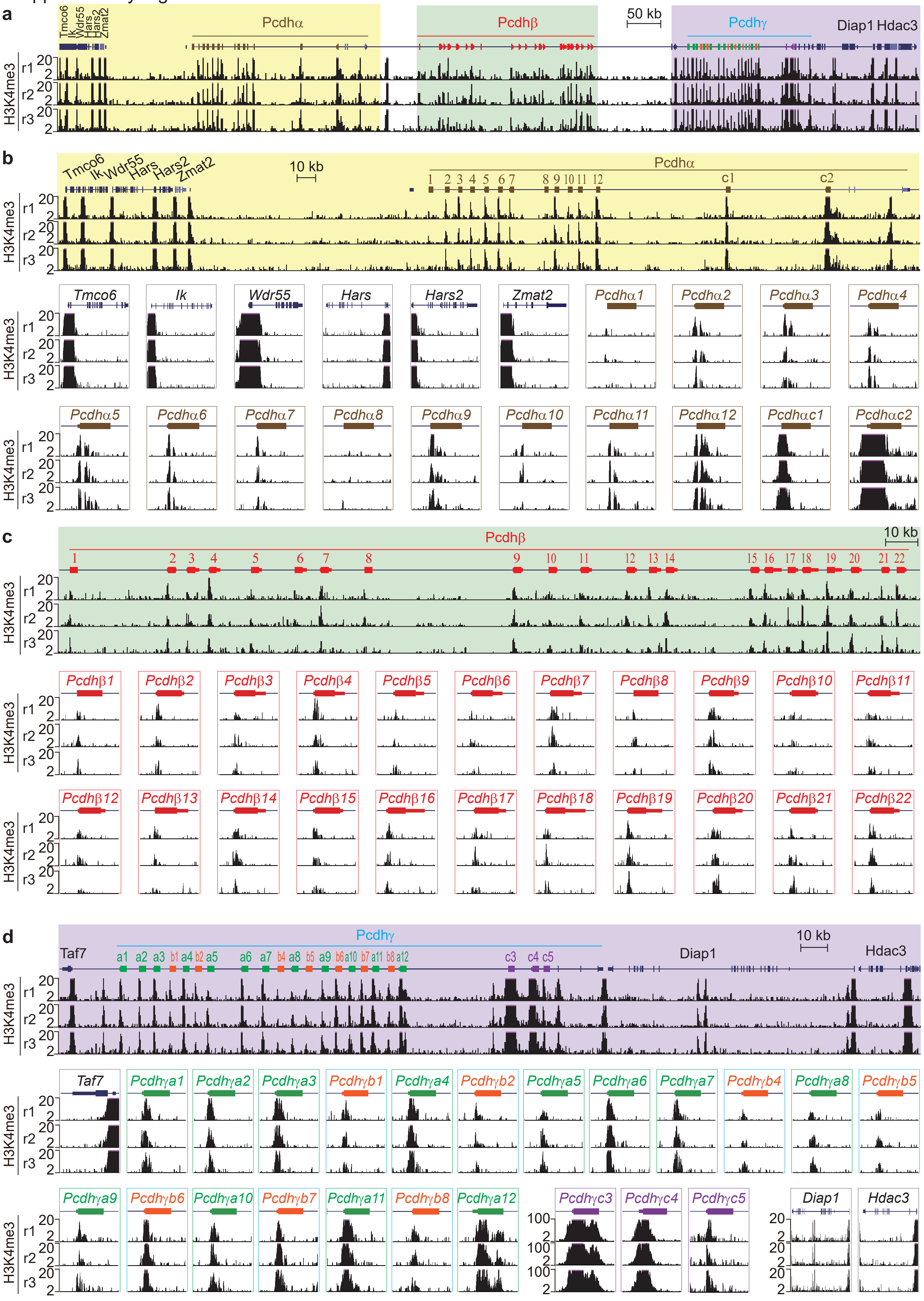

**Supplementary Fig. 3 | Enrichments of the active mark of H3K4me3 at the promoter region of each member of the three *Pcdh* gene clusters in the mouse brain. a**, H3K4me3 ChIP-seq profiles at the clustered *Pcdh* locus and its flanking regions with three replicates. **b-d,** Close-up of H3K4me3 profiles of the *Pcdh*$\alpha$ cluster (**b**), the *Pcdh*$\beta$ cluster (**c**), and the *Pcdh*$\gamma$ cluster (**d**).

Supplementary Fig4

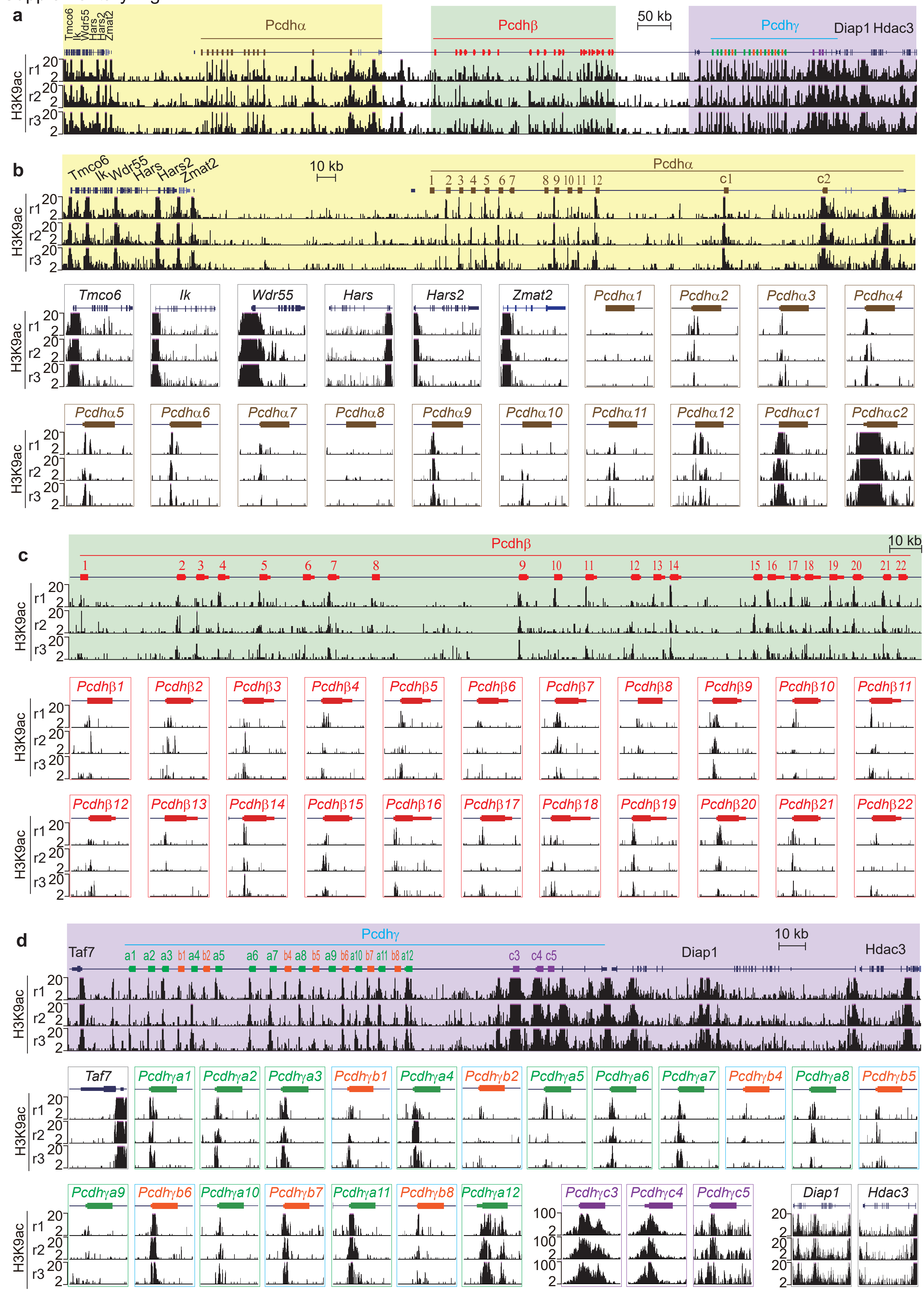

**Supplementary Fig. 4 | Enrichments of the active mark of H3K9ac at the *cPcdh* promoters and enhancers. a**, H3K9ac ChIP-seq profiles at the three *Pcdh* clusters and their flanking regions with three replicates. **b-d**, Close-up of H3K9ac profiles of the *Pcdh*$\alpha$ cluster (**b**), the *Pcdh*$\beta$ cluster (**c**), and the *Pcdh*$\gamma$ cluster (**d**).

Supplementary Fig5

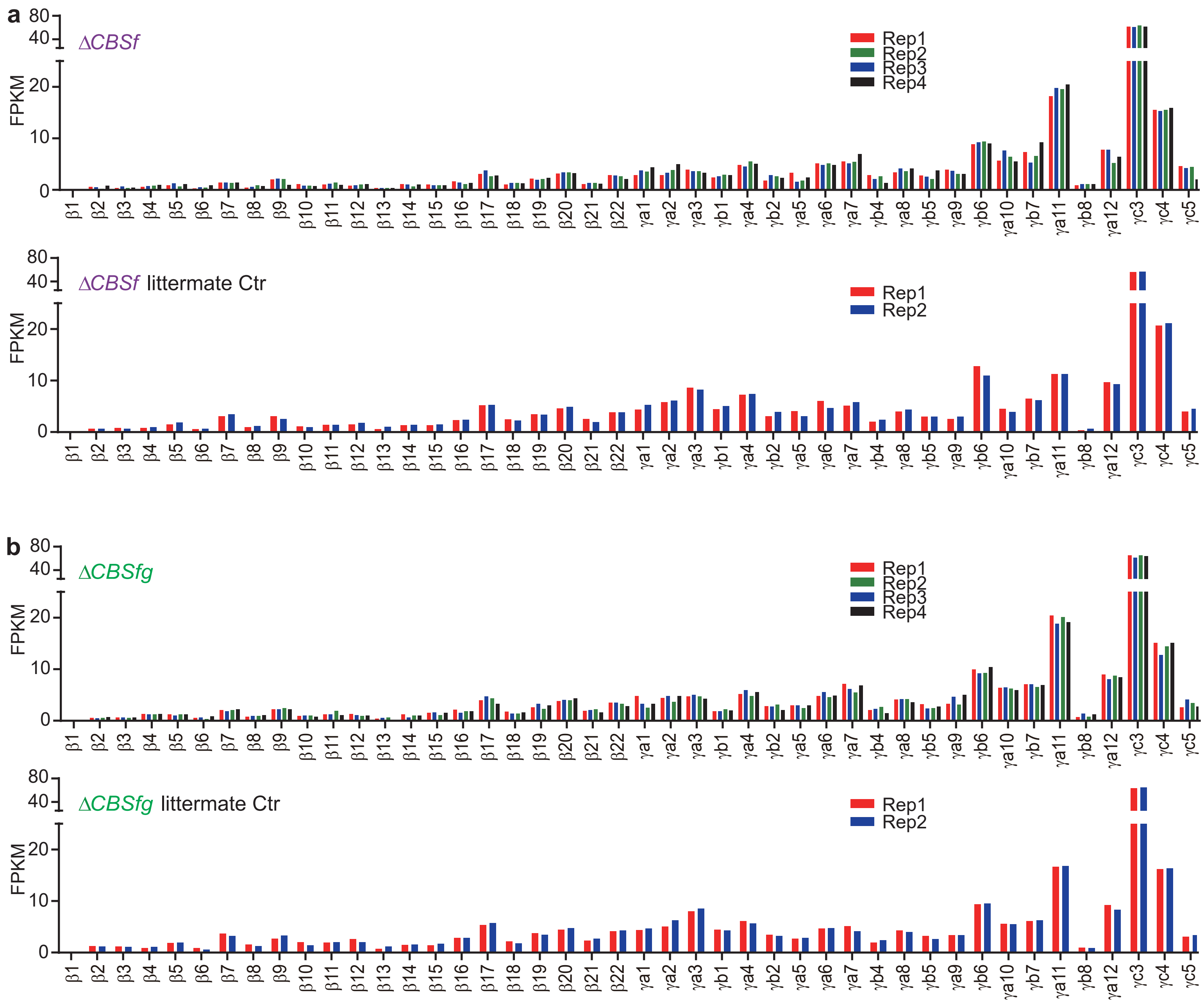

**Supplementary Fig. 5 | Replicates of RNA-seq experiments of the forward-oriented CBS-deletion mice. a-b**, Expression levels of RNA-seq replicates of microdissected neocortical tissues from the Δ*CBSf* (**a**) and Δ*CBSfg* (**b**) homozygous mice compared to their wild-type littermates. Expression levels were based on the FPKM values. Data as mean ± SD. * $P < 0.05$, ** $P < 0.01$, *** $P < 0.001$. For each mutant mouse line, two wild-type replicates and four deletion replicates were performed.

Supplementary Fig6

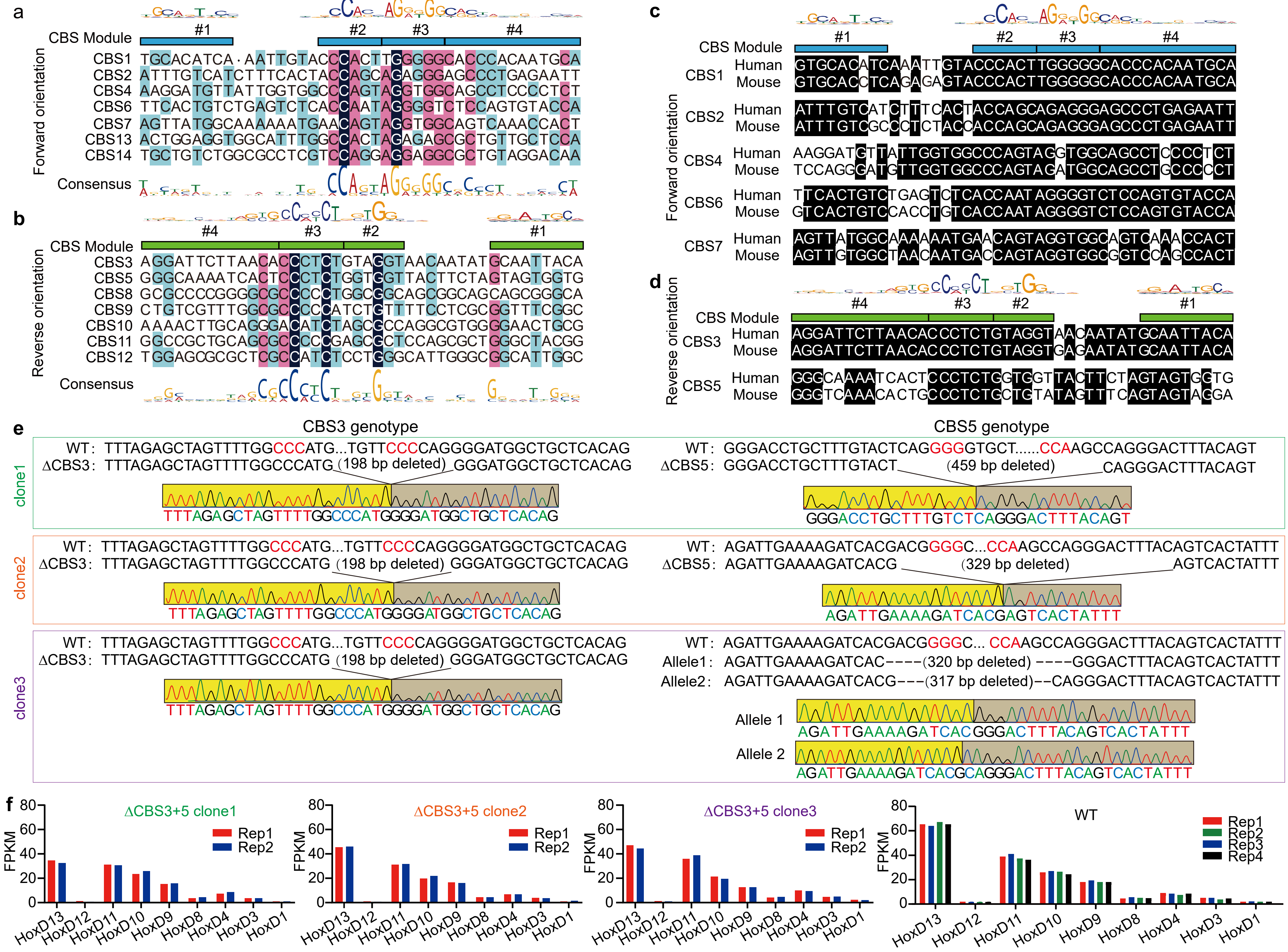

**Supplementary Fig. 6 | Decreased expression of *HOXD13* upon double knockout of outward-oriented CBS elements within the clustered CTCF TAD boundary of the HOXD gene cluster. a**,**b**, Alignments of all the forward (**a**) and reverse (**b**) CBS elements of the *HOXD* C-DOM. **c**,**d**, Conservation of the CBS elements of the *HOXD* C-DOM between human and mouse. **e**, Genotyping of *CBS3* and *CBS5* double knockout (Δ*CBS3+5*) single-cell clones by Sanger sequencing. **f**, Replicates of RNA-seq experiments showing expression levels of the *HOXD* genes in wild-type (WT) and Δ*CBS3+5* single-cell clones. Expression levels were based on the FPKM values. Data as mean ± SD. * $P < 0.05$, ** $P < 0.01$, *** $P < 0.001$.
